# Evaluating the patterning cascade model of tooth morphogenesis in the human lower mixed and permanent dentition

**DOI:** 10.1101/2023.10.13.562267

**Authors:** Dori E. Kenessey, Christopher M. Stojanowski, Kathleen S. Paul

## Abstract

**Objective:** The patterning cascade model of crown morphogenesis has been studied extensively in a variety of organisms to elucidate the evolutionary history surrounding postcanine tooth form. The current research examines the degree to which model expectations are reflected in the crown configuration of lower deciduous and permanent molars in a modern human sample. This study has two main goals: 1) to determine if metameric and antimeric pairs significantly differ in size, accessory trait expression, and relative intercusp spacing, and 2) to establish if the relative distance among early-forming cusps accounts for observed variation in accessory cusp expression.

**Methods:** Tooth size, intercusp distance, and morphological trait expression data were collected from 3D scans of mandibular dental casts representing 124 individual participants of the Harvard Solomon Islands Project. Paired tests were utilized to compare tooth size, accessory trait expression, and relative intercusp distance between diphyodont metameres and permanent antimeres. Proportional odds logistic regression was implemented to investigate how the likelihood of accessory cusp formation varies as a function of the distance between early-developing cusps.

**Results/Significance:** For paired molars, results indicated significant discrepancies in tooth size and cusp 5 expression, but not cusp 6 and cusp 7 expression. Several relative intercusp distances emerged as important predictors of accessory cusp expression. These findings support previous quantitative genetic results and suggest the development of neighboring crown structures represents a zero-sum partitioning of cellular territory and resources. As such, this study contributes to a better understanding of the evolution of deciduous and permanent molar crown configuration in humans.

## Introduction

Dental anthropological analyses often utilize nonmetric crown traits to assess the degree of similarity among global populations [1–17], detect biological kinship within a site [18–29], establish phylogenetic relationships across taxa [30–49], explore the population history associated with specific groups or geographic regions [50–70], and more recently, to estimate population affinity in recently deceased individuals as part of the biological profile [71–82]. The nonmetric crown traits that dental anthropologists rely upon to conduct these analyses include the presence and expression of accessory cusps. Accessory cusps are found in the posterior dentition (*i.e.*, premolars and molars) and begin to form after initiation of primary cusp formation [83–85]. In human molars, the early-developing primary cusps are the a) paracone, protocone, and metacone (upper molars), and b) protoconid, metaconid, hypoconid, and entoconid (lower molars) [84–91]. Late-developing upper molar accessory cusps may include the hypocone, Carabelli’s trait, cusp 5 (*i.e.*, distal accessory tubercle), and less frequently, the parastyle (*i.e.*, paramolar tubercle), while lower molar secondary cusps may consist of cusp 5 (*i.e.*, hypoconulid), cusp 6 (*i.e.*, *tuberculum sextum*), cusp 7 (*i.e.*, *tuberculum intermedium*), and less frequently, the protostylid [87, 89, 92–94].

Dental morphogenesis is the crucial embryological time frame during which the shape, size, and relative position of these accessory cusps are defined. Tooth cusp position first materializes with the appearance of the primary enamel knot (PEK) during the cap stage of dental development at the tip of the developing tooth bud [95–97]. The PEK is a transient structure formed by a cluster of tightly packed epithelial cells as a result of mesenchymal BMP4 and EDAR signaling [96, 98–100]. While the PEK does not directly map out cusp shape, size, and patterning, it has a role in inducing secondary enamel knot (SEK) production, and defects in PEK shape and size have important downstream consequences for final tooth form [97, 99, 101–103]. At the end of cap stage, MSX2, BMP2, BMP4, and the cell cycle inhibitor p21 terminate PEK cell proliferation and initiate its apoptosis [98, 104–105]. Prior to PEK dissolution, this structure prompts the formation of SEKs, an event that marks the beginning of the bell stage of dental development [97, 99, 103, 106–109].

SEK placement is an indicator of future cusp tip location through aggregate activator signaling, which promotes epithelial cell differentiation into non-proliferative knot cells (*e.g.*, FGF4, SHH), and inhibitor signaling, which represses differentiation and instead stimulates mesenchymal growth in the tooth germ (*e.g.*, BMP4, WNT6) [106, 110–112]. The complex interaction between activators and inhibitors determines the rate of proliferation across different regions of the epithelium and mesenchyme. Since the stellate reticulum prevents tissue expansion toward the root, mesenchymal proliferation induced by inhibitor signaling causes a lateral expansion in the surrounding epithelial tissue of the crown [87, 106, 113–115]. Spatial differences in cell proliferation rates around SEKs eventually cause inner enamel epithelial folding due to the steady increase in the area of the dental papilla within the spatial constraints of its surrounding enamel organ [87]. This interactive signaling network among the SEKs ensures appropriate cusp arrangement culminating in a functional tooth crown. Importantly, this signaling network governs the spatial arrangement and cusp pattern exhibited in molars, resulting in species-specific crown morphology [107, 116–117].

Evolutionary developmental (evo-devo) models have been proposed to explore the evolvability of tooth form through dental development and elucidate the evolutionary history underpinning the taxonomic diversity seen in crown morphology [93, 116, 118–119]. The patterning cascade model (PCM) specifically focuses on explaining molar crown morphogenesis [116]. The PCM highlights the complex interaction across developmental pathways through signal activation and silencing that work in tandem to shape the crown, resulting in cumulative morphological differences with each iteration. This ultimately manifests as distinct morphological patterns in posterior teeth. The model posits a higher likelihood of late-developing accessory cusp presence and/or a greater degree of accessory cusp expression with: a) shorter distances between earlier-developing cusp tips (*i.e.*, each successively appearing SEK is located directly outside of the inhibition zone of preceding SEKs), b) longer periods of crown development (*i.e.*, larger crown base), and c) more cells available for successive cusp development (larger crown size) [107, 114–116, 120].

In an initial application of this model to Lake Ladoga seals, Jernvall [116] found the presence and size of accessory cusps to be related to height differences among earlier-forming cusps: crowns with smaller cusp height differences were more likely to develop accessory cusps (Figure 1). Seal molar cusps are aligned in a single mesiodistally oriented row, with each successive cusp added in a way that maintains this linear configuration. In this case, cusp height directly shapes the volume of the mesenchymal tissue into which activators and inhibitors diffuse (*i.e.*, the taller the preceding cusps, the greater the basal cusp area falling outside the zone of inhibition, permitting the formation of additional cusps) [115–116]. However, the quadrate molars of primates are characterized by much shorter, bunodont cusps and a wider occlusal surface. In these teeth, the distance among early-developing cusps (and their associated zones of inhibition) are crucial determinants of accessory cusp development. Additionally, the number of cells responsible for carrying out cusp development decreases with each cusp added to the crown, leading to diminished stability in the form of later-developing cusps [115–116, 120]. As demonstrated in comparative studies of *Tabby* mutant and wild-type mouse dentitions, the size of the developing tooth germ also plays an important role in accessory cusp morphology [101]. Tooth germs characterized by longer developmental periods will allocate more time towards overall crown growth and additional epithelial folding to create accessory cusps [121–124]. As such, teeth characterized by a relatively large crown base area (*i.e.*, molar crowns with longer developmental periods) will have a greater likelihood of developing additional cusps [125].

**Figure 1.**
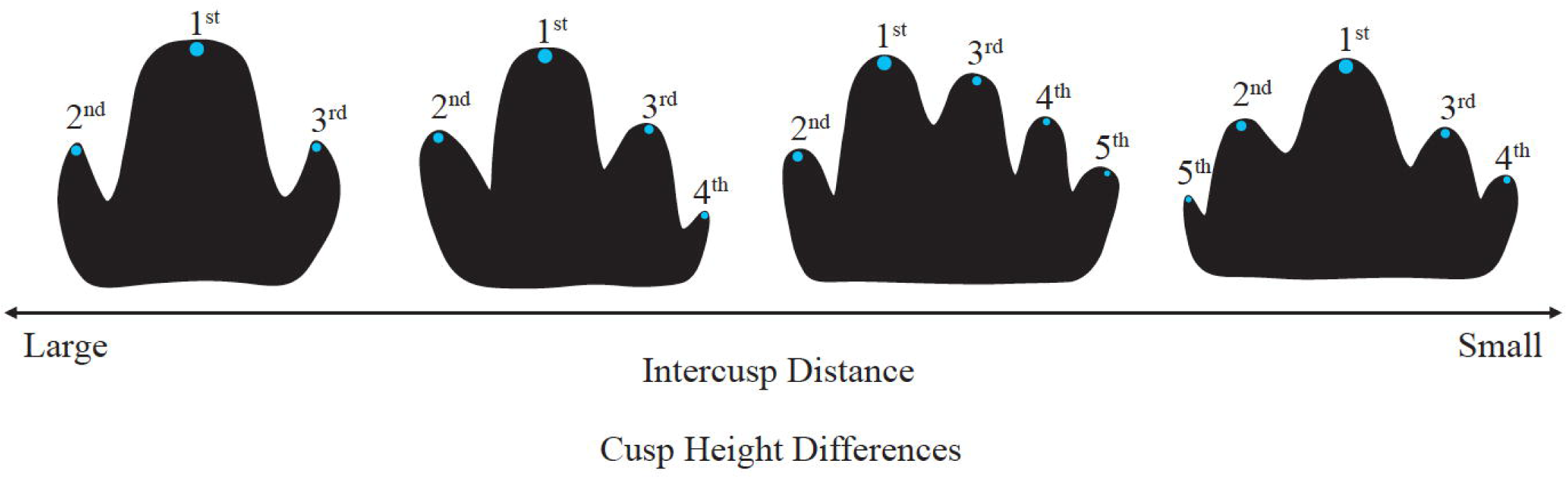
Morphological variation of Lake Ladoga seal molars following predictions of the patterning cascade model of tooth morphogenesis as outlined by Jernvall [116]. Image adapted from Jernvall & Jung [126] with permission (License number: 5642721412793). © 2000 Wiley-Liss, Inc.

The suitability of the PCM to explain mammalian molar crown morphogenesis has been tested in several taxa. Researchers found molar crown architecture to correspond to PCM expectations in Lake Ladoga seals [116, 115], mice [115], voles [115], chimpanzees [127], bonobos [127], and several human samples [128–130]. However, the PCM could only partially account for tooth form variation in American black bears [131], brown bears [131], several baboon species [132], fossil hominins [133], and other human groups [133–135]. Additionally, recent research has demonstrated that nutritional deprivation, subsistence patterns, and physiological stress may influence the measures used to test this model and the degree to which crown configuration subscribes to PCM expectations [136].

The current study expands on previous PCM research in recent modern humans by evaluating model performance in the lower diphyodont dentition. Here, we test for significant differences in tooth size, accessory cusp expression, and relative intercusp distance (RICD) between metameres (deciduous lower second molars-*dm_2_* and permanent lower first molars-*M_1_*) and antimeres (permanent left-*LM_1_* and right-*RM_1_* first molars) (Figure 2). We expect M_1_ to be significantly larger than dm_2_ [137–139], but anticipate no significant differences in size between permanent antimeres. Any significant size differences between antimeres would likely be due to fluctuating asymmetry [140–141] or random measurement error [128]. Once intercusp distances are controlled for overall crown size, we expect no significant differences in relative intercusp distances (RICDs) between metameres or antimeres. If tooth size is directly related to accessory cusp development as the PCM predicts, we anticipate more pronounced morphological trait expression in the permanent molar compared to the deciduous molar due to the expected size differences, but no significant difference between the permanent antimeres is expected.

**Figure 2.**
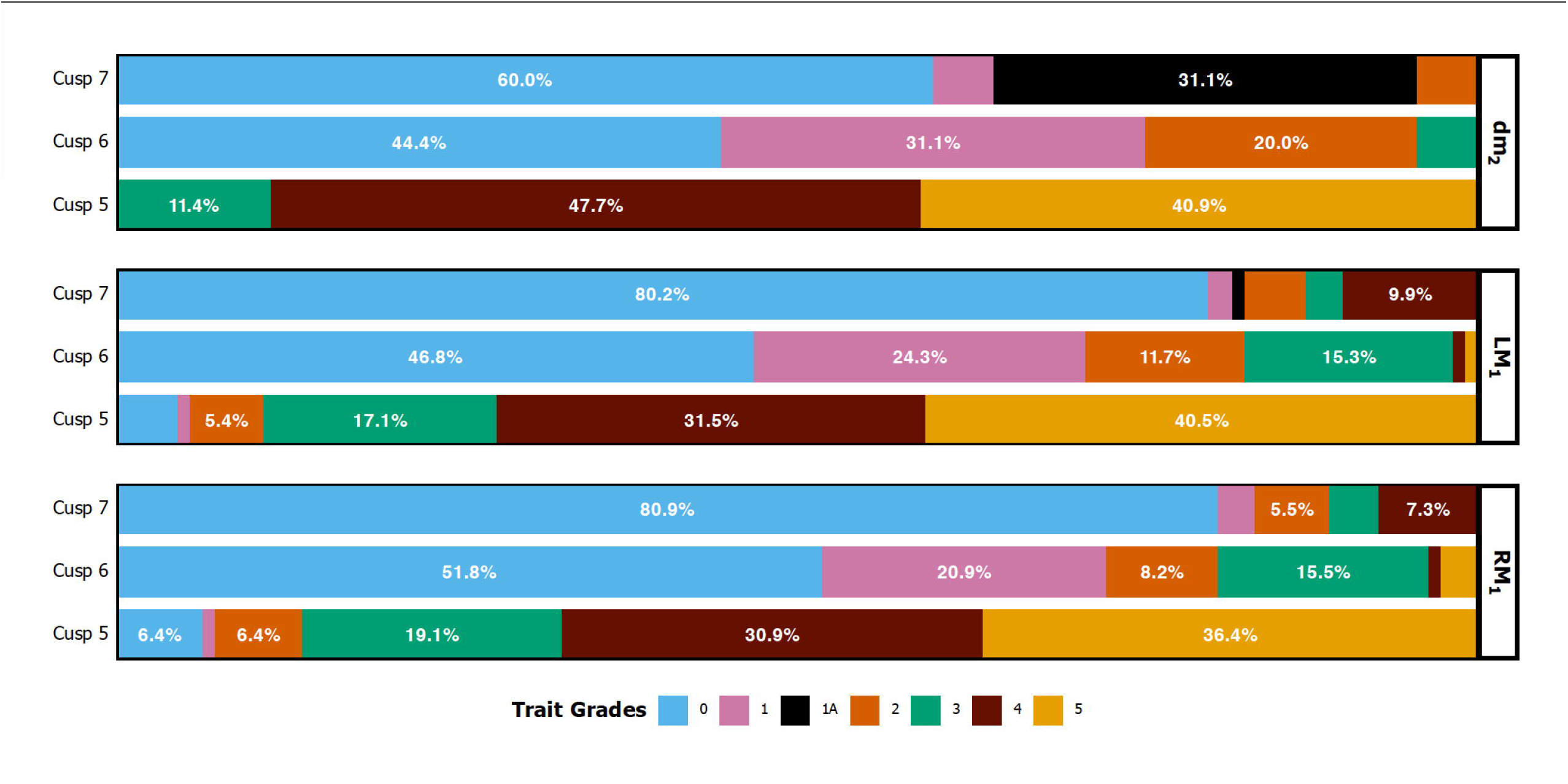
Lower molar accessory cusp trait frequencies in the study sample.

Next, we investigate whether RICDs account for variation in cusp 5 (hypoconulid), cusp 6, and cusp 7 expression. PCM predictions differ slightly for accessory cusps that are located on the occlusal surface (i.e., cusp 5, cusp 6, cusp 7) rather than on the periphery of (i.e., protosylid) human quadrate lower molar crowns to account for the zones of inhibition surrounding primary cusps [127]. These expectations are outlined in Figure 3. Following PCM predictions, we expect accessory cusps to form and exhibit higher degrees of expression if the developmental time frame of the tooth germ is relatively long and hypoconid-entoconid (cusp 5), entoconid-hypoconulid (cusp 6), and metaconid-entoconid (cusp 7) distance is expanded, thus creating sufficient space outside of inhibitory fields for an accessory cusp to form. When cusp 6 is present, we expect to see an accommodating mesiobuccal shift in cusp 5 (hypoconulid), such that the metaconid-hypoconulid distance will increase, but protoconid-hypoconulid and hypoconid-hypoconulid distance will decrease. When cusp 7 is present, we expect to see a distolingual shift in the entoconid, such that the protoconid-entoconid and hypoconid-entoconid distance will increase.

**Figure 3.**
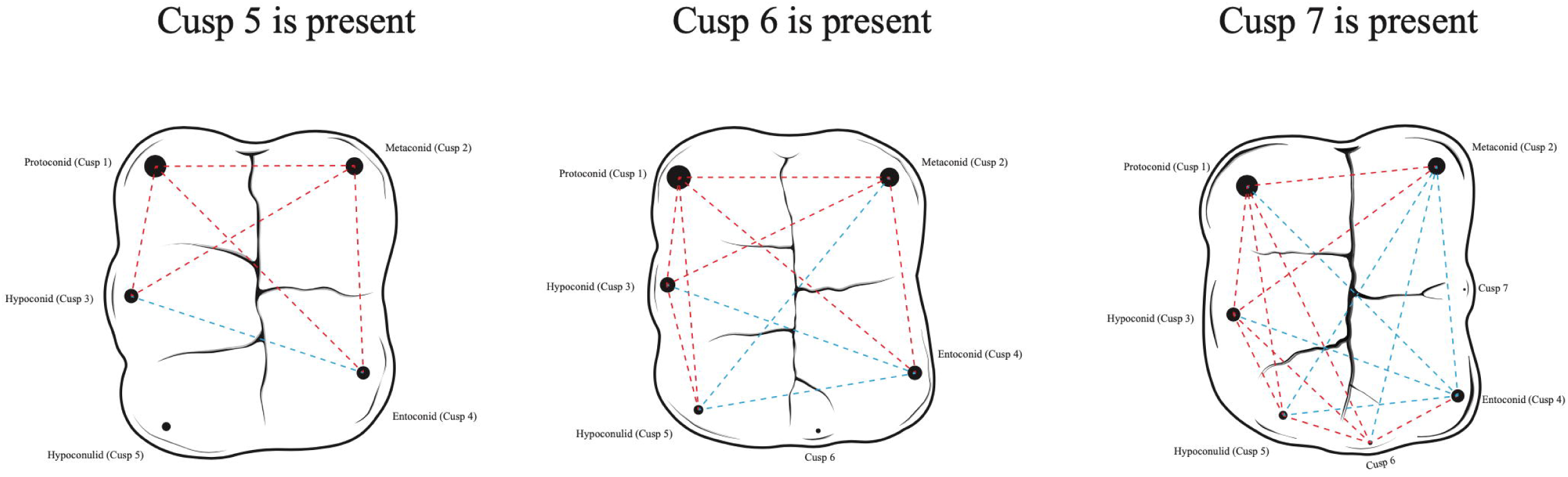
Intercusp distance expectations among early-developing cusps when a late-developing accessory cusps is present. Circles mark cusp tip position and the size of the circle corresponds to sequence of cusp development (the larger the circle, the earlier the cusp develops). Distances marked by a red dashed line indicate intercusp distances expected to be shorter when the accessory cusp is present. Distances shown with a dashed blue line illustrate intercusp distances expected to be longer when the accessory cusp is present.

## Results

Inter- and intra-observer analyses for the morphological (Table S2) and metric (Table S3) data showed strong agreement both within and between observers and are presented in the supplemental materials. To provide a succinct overview of study findings, we limit presentation of results to comparative analyses and proportional odds regression results using BLxMD crown area and RICD scaled by BLxMD area, since tooth length and width have previously been identified as important predictors of accessory cusp expression [123–124, 142–143]. All other results are presented in the S1 Appendix.

### Comparative analyses

The results of metameric and antimeric comparative analyses are presented in Table 1 for BLxMD tooth size, cusp expression, and RICD (intercusp distance scaled by BLxMD area), all other comparative results can be found in S4 Table. Differences in crown base area (p<0.00) and square root crown base area (p<0.00) were significant for both metameres (dm_2_-M_1_) and permanent antimeres (LM_1_-RM_1_). In comparing morphology, only differences in cusp 5 expression reached significance for metamere (p<0.00) and antimere (p<0.00) pairs. When examining metameric differences in paired RICD, all yielded significant differences, except distances originating from cusp 7 and the distance between cusps 1 and 5, cusps 2 and 5, and cusps 2 and 6. Between permanent antimeres, only the distance between cusps 1 and 2, cusps 2 and 7, cusps 3 and 6, and cusps 4 and 7 differed significantly. The results involving cusp 7 should be interpreted with caution as the sample of dentitions expressing cusp 7 is small (dm_2_ n=4; M_1_ n=5).

**Table 1.** Results of metameric and antimeric comparative analyses for BLxMD crown area, ASUDAS morphology, and RICD.

|  |  | Metameric Comparison |  |  |  |  | Antimeric Comparison |  |  |  |  |
| --- | --- | --- | --- | --- | --- | --- | --- | --- | --- | --- | --- |
|  |  | dm<sub>2</sub> Mean | M<sub>1</sub> Mean | rTEM (%)<sup>2</sup> | V<sup>3</sup> | p-value<sup>4</sup> | LM<sub>1</sub> Mean | RM<sub>1</sub> Mean | rTEM (%) | V | p-value |
| BLxMD Area (mm<sup>2</sup>) | Base Crown Area | **87.83** | **109.88** | **18.02** | **17** | **0.00** | **111.02** | **113.77** | **5.97** | **1983** | **0.00** |
|  | Square Root Crown Area | **9.36** | **10.46** | **8.99** | **17** | **0.00** | **10.52** | **10.65** | **3.09** | **1980** | **0.00** |
| ASUDAS | Cusp 5 | **4.31** | **3.31** | **33.65** | **480.5** | **0.00** | **3.93** | **3.79** | **9.12** | **161.5** | **0.00** |
|  | Cusp 6 | 0.84 | 1.23 | 86.55 | 76 | 0.05 | 1.00 | 0.98 | 48.33 | 121 | 0.86 |
|  | Cusp 7 | 0.2 | 0.37 | 245.37 | 4 | 0.41 | 0.60 | 0.55 | 64.92 | 50.5 | 0.36 |
| Relative Intercusp Distance (mm)<sup>1</sup> | Cusp 1-2 | **0.48** | **0.57** | **14.89** | **43** | **0.00** | **0.56** | **0.58** | **6.69** | **1681** | **0.00** |
|  | Cusp 1-3 | **0.46** | **0.51** | **13.29** | **173** | **0.00** | 0.50 | 0.49 | 8.93 | 3564 | 0.09 |
|  | Cusp 1-4 | **0.86** | **0.89** | **6.17** | **174** | **0.00** | 0.90 | 0.90 | 3.50 | 2678 | 0.33 |
|  | Cusp 1-5 | 0.86 | 0.83 | 7.96 | 580 | 0.05 | 0.83 | 0.82 | 4.45 | 3201 | 0.06 |
|  | Cusp 1-6 | **1.00** | **0.93** | **8.93** | **142** | **0.01** | 0.96 | 0.94 | 3.07 | 839 | 0.05 |
|  | Cusp 1-7 | 0.61 | 0.76 | 17.79 | 0 | 0.25 | 0.78 | 0.77 | 4.16 | 91 | 0.83 |
|  | Cusp 2-3 | **0.71** | **0.82** | **13.10** | **52** | **0.00** | 0.81 | 0.82 | 4.80 | 2818 | 0.59 |
|  | Cusp 2-4 | **0.62** | **0.65** | **6.81** | **232** | **0.00** | 0.65 | 0.66 | 5.41 | 2640 | 0.28 |
|  | Cusp 2-5 | 0.95 | 0.96 | 4.60 | 336 | 0.23 | 0.97 | 0.97 | 3.17 | 2120 | 0.09 |
|  | Cusp 2-6 | 0.89 | 0.90 | 5.86 | 62 | 0.32 | 0.91 | 0.91 | 3.34 | 553 | 0.42 |
|  | Cusp 2-7 | 0.28 | 0.35 | 30.43 | 2 | 0.75 | **0.39** | **0.37** | **9.22** | **134** | **0.03** |
|  | Cusp 3-4 | **0.70** | **0.72** | **6.05** | **293** | **0.03** | 0.72 | 0.71 | 4.41 | 3309 | 0.35 |
|  | Cusp 3-5 | **0.43** | **0.37** | **17.32** | **748** | **0.00** | 0.38 | 0.37 | 10.28 | 2867 | 0.42 |
|  | Cusp 3-6 | **0.67** | **0.56** | **15.40** | **167** | **0.00** | **0.61** | **0.57** | **15.05** | **1043** | **0.00** |
|  | Cusp 3-7 | 0.64 | 0.74 | 11.66 | 0 | 0.25 | 0.76 | 0.75 | 7.64 | 91 | 0.83 |
|  | Cusp 4-5 | **0.60** | **0.57** | **9.53** | **615** | **0.02** | 0.59 | 0.59 | 6.03 | 2609 | 0.95 |
|  | Cusp 4-6 | **0.35** | **0.38** | **15.36** | **39** | **0.04** | 0.36 | 0.37 | 9.26 | 436 | 0.05 |
|  | Cusp 4-7 | 0.31 | 0.31 | 36.32 | 3 | 1 | **0.32** | **0.36** | **14.33** | **39** | **0.04** |
|  | Cusp 5-6 | **0.40** | **0.30** | **24.09** | **168** | **0.00** | 0.33 | 0.31 | 17.94 | 814 | 0.09 |
|  | Cusp 5-7 | 0.76 | 0.76 | 6.37 | 3 | 1 | 0.77 | 0.77 | 4.19 | 94 | 0.43 |
|  | Cusp 6-7 | 0.69 | 0.59 | 11.25 | 1 | 1 | 0.60 | 0.66 | 8.86 | 0 | 0.13 |

<sup>1</sup>Intercusp distance scaled by BLxMD tooth size <sup>2</sup>rTEM: Relative technical error of measurement <sup>3</sup>V: Paired-samples Wilcoxon signed-rank test statistic <sup>4</sup>α≤0.05; significant results in bold

### Regression analyses

We first tested the PCM assumption that an extended period of crown development (using crown base area as a proxy) increases the likelihood of developing accessory lower molar cusps with higher trait expression (S6 Table). Tooth size was not a significant predictor of cusp 5 (p=0.82), cusp 6 (p=0.30), or cusp 7 (p=0.16) expression in dm_2_. In permanent molars, tooth size was a significant predictor of cusp 5 expression for the left (p=0.03) and the right (p=0.01) antimere, as well as cusp 6 (p=0.05) expression in RM_1_. Tooth length was an important predictor of permanent cusp 5 expression only, with the odds of increased expression elevated by 7% for LM_1_ and 11% for RM_1_ for every 0.10 unit increase in mesiodistal diameter. On the other hand, an increase in cusp 6 expression appears to be associated with tooth width, rather than length, with the odds of developing a cusp 6 with higher grade expression increasing by 8% for every 0.10 unit increase in buccolingual RM_1_ dimension. Conversely, having a longer molar crown increases the likelihood of developing a cusp 5 with greater expression, while having a wider crown boosts the probability of developing a cusp 6 with greater expression. These findings may be influenced by sex-specific differences in tooth dimensions and/or morphological trait expression.

We next examined RICD as a predictor of accessory cusp expression, scaling the intercusp distances by crown area (BLxMD) to eliminate tooth size as a confounding factor in the analyses (Figure 4-5). For dm_2_, the only significant predictor of accessory cusp expression was the distance between cusps 4 and 5, those with relatively higher intercusp distances being 9.65 times more likely to develop a cusp 6 with higher expression for every 0.10 unit increase in intercusp distance. For both permanent antimeres, a diminished distance between cusps 1 and 3 and cusps 2 and 3 increases the likelihood of developing a cusp 5 with higher expression. For LM_1_, the likelihood of developing a cusp 5 with higher expression increases by 32% and 36% for every 0.10 unit decrease in cusp 1-3 and cusp 2-3 distance, respectively. For RM_1_, the likelihood of developing a cusp 5 with higher expression increases by 25% and 40% for every 0.10 unit decrease in cusp 1-3 and cusp 2-3 distance, respectively.

**Figure 4.**
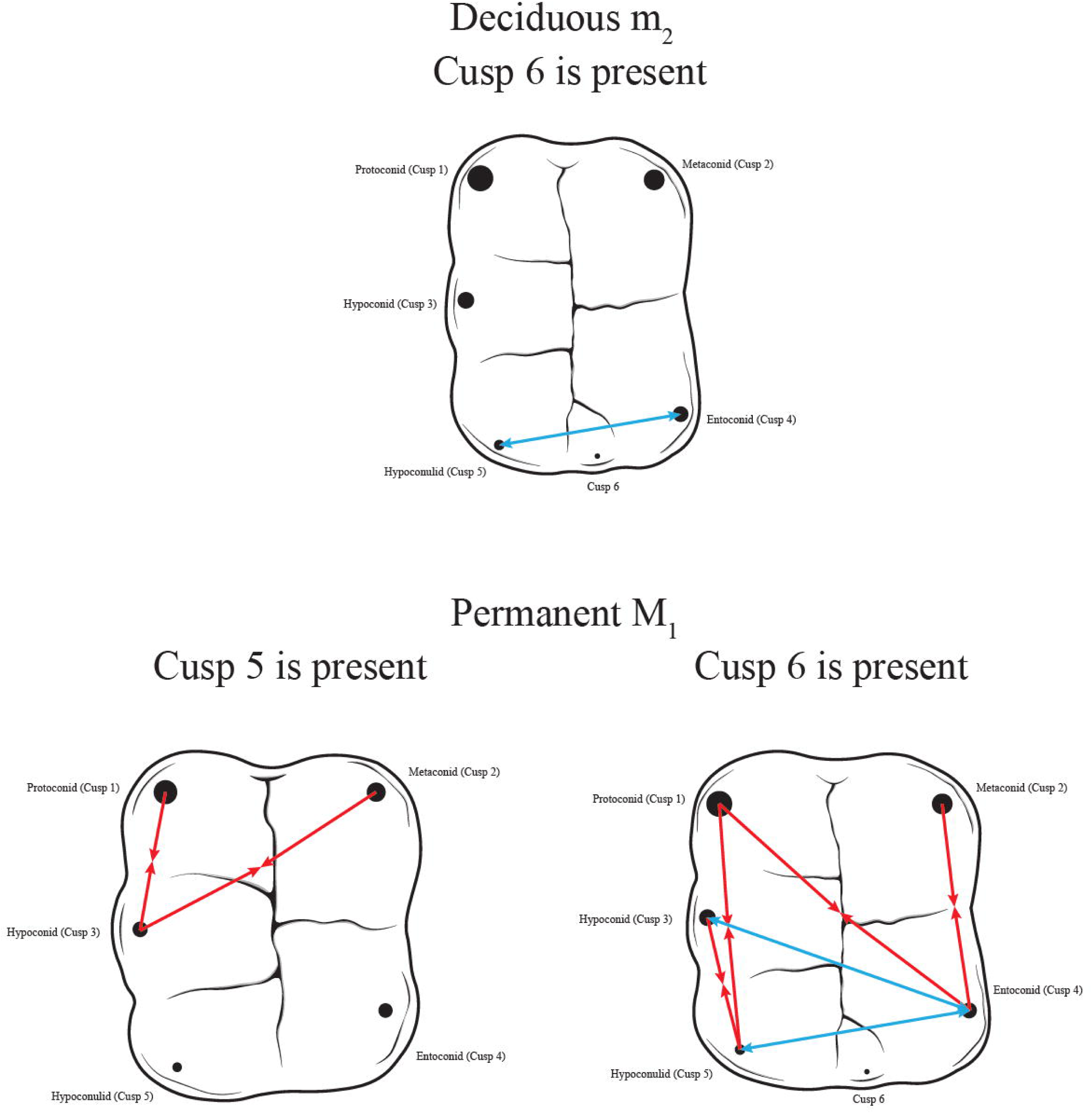
Relative intercusp distances that significantly contribute to the likelihood of accessory cusp expression in both permanent antimeres. Red arrows pointing toward each other indicate a negative relationship between intercusp distance and morphological trait expression. Blue arrows pointing away from each other indicate a positive relationship between intercusp distance and morphological trait expression.

**Figure 5.**
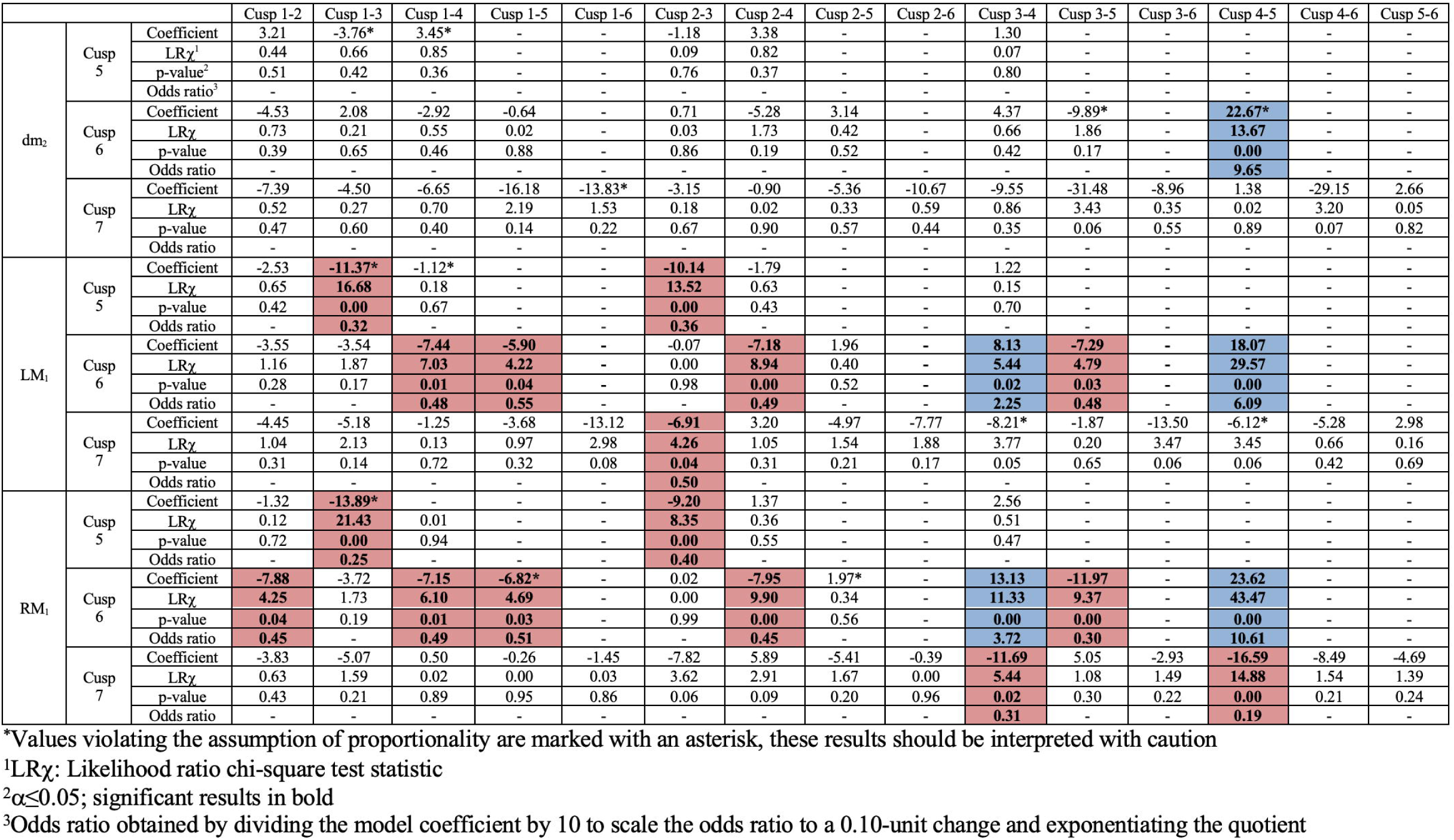
Results of proportional ordinal regression when using RICD scaled by BLxMD crown area as a predictor of lower molar accessory cusp trait expression. Cells highlighted in blue represent intercusp distances that become significantly longer with increasing accessory cusp expression. Cells highlighted in red represent intercusp distances that become significantly shorter with increasing accessory cusp expression.

Numerous intercusp distances play an important role in permanent molar cusp 6 expression. For both antimeres, a reduced distance between cusps 1 and 4, cusps 1 and 5, cusps 2 and 4, and cusps 3 and 5 increases the likelihood of developing a cusp 6 with higher expression. An increased distance between the cusps surrounding cusp 6 increases the likelihood of developing an accessory cusp with higher expression. On the right side, the chances of developing a more pronounced cusp 6 increases about four-fold and 10-fold with each additional 0.10 unit increase in cusp 3-4 and cusp 4-5 distance, respectively. On the left side, the chances of developing a cusp 6 with higher expression increases 2.25 and 6 times, for every 0.10 unit increase in the respective intercusp distances. These results are consistent with our original prediction that cusp 6 development will cause a mesiobuccal shift in the hypoconulid as evidenced by the significant role of a diminished cusp 1-5 and cusp 3-5 distance in cusp 6 development. While the predicted increase in cusp 2-5 distance is also observable, this variable did not play a significant role in cusp 6 expression as was initially expected.

Cusp 7 expression is shaped by the distance between cusps 2 and 3 on the left side. The likelihood of developing a cusp 7 with higher expression increases by 50% for every 0.10 unit decrease in the distance between cusps 2 and 3. On the right side, a reduced distance between cusps 3 and 4 as well as cusps 4 and 5 appear to be important predictors of cusp 7 expression. For every additional 0.10 unit decrease in the distance between cusps 3-4 and 4-5, there is 69% and 81% higher odds of developing a more pronounced cusp 7, respectively. The role of an increased cusp 3-4 distance seen in the right permanent antimere is consistent with the expectation of a distolingual shift in the entoconid outlined in the study hypotheses, however, this relationship only approached significance for the left antimere. Interestingly, the predicted distance increase between cusps 2-4 to accommodate the accessory cusp wedged between these two primary cusps was also not a significant predictor of trait expression.

## Discussion

### Comparative Analyses

One of the goals of the current study was to compare morphological trait expression, crown size, and intercusp distance in paired molars. This study identified significant differences in tooth size and cusp 5 expression between deciduous and permanent metameres, corroborating previous findings [134]. The significant difference in metameric cusp 5 expression may be due to disparate susceptibility to environmental influences characterizing the deciduous and permanent dentitions. The dm_2_ is very likely to exhibit a cusp 5, while cusp 5 expression on M_1_s tends to be more variable. This could be consistent with the idea that the deciduous dentition is more “canalized” and less susceptible to environmental influence than its permanent counterpart [27, 144–145], though recent heritability estimate comparisons between metameres failed to find support for this hypothesis [146]. Significant difference in cusp 5 expression between antimeres may also support the idea that this cusp is susceptible to environmental influences, which is consistent with previous findings [141, 147]. Non-significant differences in metameric and antimeric cusp 6 expression are consistent with previous research, but the lack of difference in cusp 7 expression between tooth pairs diverges from other studies, which identified more pronounced cusp 7 expression in dm_2_ [148–149]. The lack of significant difference in metameric and antimeric cusp 6 and cusp 7 expression may indicate that these accessory cusps are more resilient to environmental disturbances than cusp 5 development. This rationale is bolstered by the findings of heritability studies, which have identified cusp 6 and cusp 7 to be under strong genetic control [146, 150–151]. This result may also reflect the outcome of inhibitory zoning when cusp 5 and cusp 6 are expressed concurrently on the distal crown. There is likely more room for variation in cusp 5 expression when there are no other cusps competing for cellular resources on the distal aspect of the crown.

Paired tests indicate that most RICDs significantly differ between metameres. For distances involving cusp 7, inconsistent results for the antimeric and metameric comparisons may relate to the relatively small number of individuals that possess this accessory cusp within the study sample. Differences between the other RICDs may reflect the distinct crown configurations between deciduous and permanent elements, in particular, significant discrepancies in cusp 5 expression (see Table 1). The reason for significantly different cusp 1-2 and cusp 3-6 distances between the permanent antimeres is also unclear. It is possible however, that because these dimensions involved the earliest-forming cusps, they have the greatest potential for variation. Beyond sample size concerns, it is likely that the non-significant RICD differences between metameres and the significant differences in the RICD differences between antimeres are caused by random measurement error [128].

### Regression analyses

Another important aim of this study was to establish if tooth size and RICDs, following the PCM, were important predictors of accessory cusp formation. This study is the first to test PCM expectations in lower dm_2_, though previous work has explored this model and its associated predictions for Carabelli’s trait in dm^2^ [134]. Overall neither tooth size nor RICD seems to be a predictor of deciduous lower molar accessory cusp expression. The only significant predictor of cusp 6 expression was found to be an increased distance between cusps 4 and 5, which is consistent with PCM expectations. The overall absence of a relationship between dm_2_ RICD, tooth size, and accessory cusp expression differs from the findings of Paul and colleagues [134], who identified more consistent relationships between RICDs and accessory cusp expression in the deciduous rather than permanent dentition. The current results may diverge from those of Paul and colleagues [134] due to the lack of variation exhibited by cusp 5 and the limited number of individuals exhibiting cusp 7. Another plausible explanation may be the differing cusp configurations that characterize maxillary and mandibular molars. The peripheral location of Carabelli’s trait may correspond to PCM expectations more closely than the occlusally located accessory cusps examined in this study. Since no other study has tested PCM predictions in dm_2_, further research with larger sample sizes, samples characterized by a wider range of morphological variation, and samples coming from diverse populations would help contextualize the current results. Nevertheless, the ability of the current study to highlight the importance of the distance between cusps 4 and 5 in predicting cusp 6 expression indicates that cusp 6 is a true accessory cusp with a developmental pattern closely shaped by the spatial relationship among earlier-developing cusps.

Size-related variables were significant predictors of accessory cusp expression in permanent molars, particularly tooth length for cusp 5 and tooth width for cusp 6 and 7. These results align with those of previous work that identified larger tooth size as a predictor of increased crown complexity [121, 123–124, 127, 142]. In particular, Dahlberg [121] and Garn and colleagues [142], found longer mesiodistal M_1_ dimension to increase the likelihood of developing a five-cusped tooth. To our knowledge, the relationship between buccolingual dimension and cusp 6 and/or 7 development has not been directly examined, though Skinner and Gunz [127] did find that members of the genus *Pan* with a cusp 6 had significantly larger teeth. A buccolingual expansion of the crown would increase the likelihood that areas along the mesial and distal marginal ridge fall outside the inhibition zones of primary cusps, thus promoting accessory cusp development following the PCM.

For permanent antimeres, a reduced protoconid-hypoconid and metaconid-hypoconid distance increased the likelihood of developing higher cusp 5 expression. These findings are consistent with PCM expectations, whereby a mesiolingual shift in the hypoconid would serve to provide adequate space for the development of a cusp 5 that is more proximate to the hypoconid than the entoconid. While an increased hypoconid-entoconid distance was observed in both antimeres following the PCM, none of these variables reached the level of significance. The lack of such a relationship in the current study differs from the findings of Ortiz and colleagues [133], who observed a significant increase in hypoconid-entoconid distance with cusp 5 development in both fossil hominins and recent modern humas. This study’s failure to replicate previous results may relate to divergent sample composition. It is also possible that a transverse expansion of the talonid is not strictly necessary to accommodate the development of cusp 5 since the hypoconid and entoconid may be sufficiently far apart already in a four-cusped molar. This explanation is also supported by the findings that a buccolingual expansion in lower permanent molars does not increase the likelihood of developing a cusp 5 with more pronounced expression.

Several RICD variables emerged as important predictors of cusp 6 expression. Consistent with our initial predictions, the results of the proportional odds logistic regression indicate a mesiobuccal shift in cusp 5 (hypoconulid) to integrate cusp 6. However, a mesiobuccal shift in the entoconid is also revealed by a reduced protoconid-entoconid and metaconid-entoconid distance in individuals with more pronounced cusp 6 expression. This mesiobuccal shift seen in the entoconid is significant enough that the protoconid-entoconid distance is reduced, but not excessively, as increased hypoconid-entoconid distance predicts larger cusp 6 expression. Supporting the current study’s original hypothesis, an increased entoconid-hypoconulid distance was a significant predictor of cusp 6 expression. The relationship between entoconid-hypoconulid distance and cusp 6 appears to be a crucial predictor of the expression displayed by this accessory cusp since its role in cusp 6 expression has been replicated in the *Pan*, *Pongo*, *Australopithecus*, *Paranthropus*, and *Homo* genera, as well as another recent modern human sample [127, 133]. This suggests that while a reduction in intercusp distances is an important predictor of accessory cusp development, increasing the distance between certain cusps to expose regions of the crown outside of inhibition zones may be equally important for accessory cusp development, which is not explicitly outlined in the PCM [116]. The importance of expanded distances between certain earlier developing cusps is more than likely related to the integrated cuspal arrangement characterizing the quadrate molars of primates. Accessory cusps developing on the primate posterior dentition are more occlusally located on the tooth crown as opposed to a peripheral position typical of accessory cusps on Lake Ladoga seal molars—the elements to which the PCM was first applied.

The antagonistic relationship between cusp 5 and cusp 6 expression revealed by quantitative genetic studies may also explain why these accessory cusps subscribe to PCM expectations to different degrees. Stojanowski and colleagues [147] and Paul and colleagues [152] uncovered a negative genetic correlation between cusp 5 and cusp 6 development in lower molars, indicating that these cusps may be competing for “cellular real estate” during development and the amount of resources allotted to one of these accessory cusps will have direct and opposing consequences on the development and expression of neighboring accessory cusps. In concert, these findings indicate that while cusp 6 expression may follow PCM expectations closely, the development of this cusp will cause more variability in cusp 5 expression, likely leading to deviations in the degree to which this accessory cusp conforms to model expectations.

This study was unable to identify a significant relationship between increased metaconid-entoconid distance and cusp 7 expression as was hypothesized prior to analysis. The struggle to identify RICDs as consistent predictors of cusp 7 development is not unique to the current project. Ortiz and colleagues [133] were also unable to detect a relationship between metaconid-entoconid distance and cusp 7 expression in the *Australopithecus*, *Homo*, and *Pan* genera. Kozitzky [132] sought to determine if the expression of the median lingual notch cuspule (MLNC) in the upper molars of several baboon groups follows PCM predictions. Based on its location, the MLNC may be an isomeric structure in the upper molars of *Cercopithecoidea* that is similar to cusp 7, although its spatial placement indicates a much closer association with the cingulum than cusp 7, which is more closely integrated within the crown. In a pooled sample, protocone-hypocone distance was not a significant predictor of MLNC expression, but when considering only the first molar, this variable was significant in hamadryas baboons [132]. The failure of cusp 7 expression to follow PCM expectations across several studies, may indicate that a different developmental pathway may shape cusp 7 development [153]. It is also possible that the overall size variation characterizing cusp 7 expression is fairly minimal and the trait expression pattern may be difficult to detect on the outer enamel surface versus the enamel-dentine junction due to factors like enamel thickness or the presence of a corresponding dentine horn. This variation can lead to conflation of the “metaconulid-type” and “interconulid-type” cusp 7 in analyses that rely on trait expression of the external crown surface exclusively [153–157].

In sum, the current study found partial support for the PCM in dm_2_ and M_1_. Similar to previous studies, findings consistent with PCM expectations involve cusp 6 [127, 133]. Human upper molars have received more attention in PCM studies, with some outlining cusp patterns consistent with the PCM [117, 128] and others only identifying partial agreement with the model [133–135]. The inability of the PCM to consistently account for postcanine cusp development, size, and arrangement potentially indicates that this model alone may not be sufficient to explain the morphological diversity characterizing the human dentition. It is possible that certain model organisms with less derived dental patterns, like Lake Ladoga seals on which the model was developed, conform more closely to PCM expectations due to more canalization in tooth form and less susceptibility to external factors shaping development. While Salazar-Ciudad and Jernvall [114] were able to reconstruct the general quadritubercular form that resembles human upper molars using PCM simulations, they acknowledge that model parameters (*e.g.*, epithelial growth rate, rate of epithelial activator secretion) can be modified by population variation and environmental components, though the degree to which crown form can deviate from the expected pattern due to these factors is not well-understood. The extent to which environmental variables can infiltrate cusp morphogenesis is relatively understudied, but recent research suggests nutritional deprivation and subsistence strategy can alter accessory trait expression, crown size, and intercusp spacing [136]. Following Salazar-Ciudad and Jernvall [114], it is possible that environmental factors may be the reason for inconsistencies. Additional research expanding the sample composition (*e.g.*, incorporating populations of diverse bioregional affiliation), the variables examined (*e.g.*, deciduous and permanent antimeres, metameres, and isomeres), and accounting for various environmental factors (*e.g.*, intrauterine environment, nutritional deprivation, pathological conditions, physiological stress) in future studies will shed light on how and to what extent these elements alter cusp arrangement compatibility with PCM predictions.

## Materials and Methods

A detailed overview of study materials and methods is presented in the S2 Appendix. The study examined tooth size and morphological trait expression for 124 individuals using 3D digital scans of the lower dentition (see Figure 2). Tooth size dimensions and morphological trait expression were assessed using established standards in the field [158–161]. Paired sample tests were used to detect significant differences in tooth size and morphological trait expression between diphyodont metameres and permanent antimeres. Relative intercusp distances were established by scaling intercusp dimensions by overall crown size to eliminate it as a confounding variable. Proportional odds regression was utilized to test if the distances between early-forming cusps were significant predictors of lower molar accessory cusp expression.

## Acknowledgements

This research was funded by the National Science Foundation (BCS-1540313; BCS-1750089; BCS-1063942) and the Wenner–Gren Foundation (Dissertation Fieldwork Grant). We would like to thank Dr. Andrew Seidel for helping to generate the images and scans used in this study. We would like to acknowledge the critical support provided by the staff of the Peabody Museum of Archaeology and Ethnology, Harvard University for this project.

## Supporting Information

**S1 Appendix. Supplementary figures and tables**

**S2 Appendix. Extended description of materials and methods**

