## Supplementary material for "Evaluating the patterning cascade model of tooth morphogenesis in the human lower mixed and permanent dentition": S1 Appendix

**Supplement Glossary**

- AICA: absolute intercusp area
- BL: buccolingual tooth dimension
- BLxMD: crown area obtained by multiplying tooth length (MD) by tooth width (BL)
- BLICA: relative intercusp area scaled by buccolingual tooth dimension
- Cusp 1-2: Intercusp distance between protoconid and metaconid
- Cusp 1-3: Intercusp distance between protoconid and hypoconid
- Cusp 1-4: Intercusp distance between protoconid and entoconid
- Cusp 1-5: Intercusp distance between protoconid and hypoconulid (cusp 5)
- Cusp 1-6: Intercusp distance between protoconid and cusp 6
- Cusp 1-7: Intercusp distance between protoconid and cusp 7
- Cusp 2-3: Intercusp distance between metaconid and hypoconid
- Cusp 2-4: Intercusp distance between metaconid and entoconid
- Cusp 2-5: Intercusp distance between metaconid and hypoconulid (cusp 5)
- Cusp 2-6: Intercusp distance between metaconid and cusp 6
- Cusp 2-7: Intercusp distance between metaconid and cusp 7
- Cusp 3-4: Intercusp distance between hypoconid and entoconid
- Cusp 3-5: Intercusp distance between hypoconid and hypoconulid (cusp 5)
- Cusp 3-6: Intercusp distance between hypoconid and cusp 6
- Cusp 3-7: Intercusp distance between hypoconid and cusp 7
- Cusp 4-5: Intercusp distance between entoconid and hypoconulid (cusp 5)
- Cusp 4-6: Intercusp distance between entoconid and cusp 6
- Cusp 4-7: Intercusp distance between entoconid and cusp 7
- Cusp 5-6: Intercusp distance between hypoconulid (cusp 5) and cusp 6
- Cusp 5-7: Intercusp distance between hypoconulid (cusp 5) and cusp 7
- Cusp 6-7: Intercusp distance between cusp 6 and cusp 7
- MAE: mean absolute error
- MAPE: mean absolute percentage error
- MD: mesiodistal tooth dimension
- MDICA: relative intercusp area scaled by mesiodistal tooth dimension
- RICA: relative intercusp area scaled by BLxMD crown area
- RMSE: root-mean-square error
- rTEM: relative technical error of measurement
- TEM: technical error of measurement
- TICA: relative intercusp area scaled by traced crown area
- Trace: crown area obtained by tracing the crown outline

**S1 Table. Grading system used to assess accessory cusp expression following Scott & Irish [160] and grade frequencies for study sample**

| Trait | Grade | Definition | dm_2_ | | LM_1_ | | RM_1_ | |
| --- | --- | --- | --- | --- | --- | --- | --- | --- |
|  |  |  | % | n | % | n | % | n |
| Hypoconulid (Cusp 5) | 0 | Hypoconulid is absent (four-cusped tooth) | 0.00 | 0/44 | 4.50 | 5/111 | 6.36 | 7/110 |
|  | 1 | Trace expression | 0.00 | 0/44 | 0.90 | 1/111 | 0.09 | 1/110 |
|  | 2 | Slight | 0.00 | 0/44 | 5.40 | 6/111 | 6.36 | 7/110 |
|  | 3 | Moderate | 11.4 | 5/44 | 17.1 | 19/111 | 19.1 | 21/110 |
|  | 4 | Strong | 47.7 | 21/44 | 31.5 | 35/111 | 30.9 | 34/110 |
|  | 5 | Pronounced | 40.9 | 18/44 | 40.5 | 45/111 | 36.4 | 40/110 |
| Cusp 6  (*Tuberculum sextum*) | 0 | Absence of cusp 6 | 44.4 | 20/45 | 46.8 | 52/111 | 51.8 | 57/110 |
|  | 1 | Cusp 5 is more than twice the size of cusp 6 | 31.1 | 14/45 | 24.3 | 27/111 | 20.9 | 23/110 |
|  | 2 | Cusp 5 is about twice the size of cusp 6 | 20.0 | 9/45 | 11.7 | 13/111 | 8.18 | 9/110 |
|  | 3 | Cusp 5 and cusp 6 are about equal in size | 4.44 | 2/45 | 15.3 | 17/111 | 15.5 | 17/110 |
|  | 4 | Cusp 6 is slightly larger than cusp 5 | 0.00 | 0/45 | 0.09 | 1/111 | 0.09 | 1/110 |
|  | 5 | Cusp 6 is markedly larger than cusp 5 | 0.00 | 0/45 | 0.09 | 1/111 | 2.73 | 3/110 |
| Cusp 7  (*Tuberculum intermedium*) | 0 | No accessory cusp between cusps 2 and 4 | 60.0 | 27/45 | 80.2 | 89/111 | 80.9 | 89/110 |
|  | 1A | This expression does not assume the typical wedge-shaped form of a cusp 7, but is marked by a groove on the lingual surface of the metaconid | 31.1 | 14/45 | 0.09 | 1/111 | 0.00 | 0/110 |
|  | 1 | Small, wedge-shaped cusp between cusps 2 and 4 | 4.44 | 2/45 | 1.80 | 2/111 | 2.73 | 3/110 |
|  | 2 | Distinct, but small cusp | 4.44 | 2/45 | 4.50 | 5/111 | 5.45 | 6/110 |
|  | 3 | Moderate cusp | 0.00 | 0/45 | 2.70 | 3/111 | 3.64 | 4/110 |
|  | 4 | Large cusp | 0.00 | 0/45 | 9.91 | 11/111 | 7.27 | 8/110 |

|  | Intraobserver error | | | Interobserver error | | |
| --- | --- | --- | --- | --- | --- | --- |
|  | Cusp 5 | Cusp 6 | Cusp 7 | Cusp 5 | Cusp 6 | Cusp 7 |
| LM_1_ | 0.913 | 0.958 | 0.853 | 0.865 | 0.866 | 0.856 |
| RM_1_ | 0.96 | 0.91 | 0.893 | 0.802 | 0.785 | 0.674 |

**S2 Table. Cohen’s weighted kappa (κ) indicating observer agreement for morphological traits**

**S3 Table. Different measures of intra-observer error for absolute intercusp distance measurements**

|  | dm_2_ | | | | | LM_1_ | | | | | RM_1_ | | | | |
| --- | --- | --- | --- | --- | --- | --- | --- | --- | --- | --- | --- | --- | --- | --- | --- |
|  | MAE | MAPE | RMSE | TEM | rTEM | MAE | MAPE | RMSE | TEM | rTEM | MAE | MAPE | RMSE | TEM | rTEM |
| Cusp 1-2 | 0.3139333 | 7.10% | 0.3681528 | 0.2603233 | 5.68% | 0.3875333 | 6.73% | 0.4341853 | 0.3070154 | 5.17% | 0.3854667 | 6.38% | 0.4881221 | 0.34515450 | 5.74% |
| Cusp 1-3 | 0.2672667 | 6.38% | 0.3187249 | 0.2253725 | 5.35% | 0.5300000 | 10.37% | 0.6163899 | 0.4358535 | 7.99% | 0.3276667 | 6.59% | 0.4005374 | 0.28322270 | 5.40% |
| Cusp 1-4 | 0.4729333 | 6.03% | 0.6026081 | 0.4261083 | 5.23% | 0.5196667 | 5.67% | 0.6527354 | 0.4615536 | 4.88% | 0.4068667 | 4.58% | 0.4919395 | 0.34785380 | 3.71% |
| Cusp 1-5 | 0.3966667 | 5.06% | 0.5282853 | 0.3735541 | 4.61% | 0.6843846 | 8.01% | 0.7882435 | 0.5573723 | 6.34% | 0.4507500 | 5.35% | 0.4981901 | 0.35227360 | 4.08% |
| Cusp 1-6 | 0.2147500 | 2.34% | 0.3153213 | 0.2229658 | 2.37% | 0.8515000 | 9.21% | 0.9422196 | 0.6662499 | 6.86% | 0.4504000 | 4.61% | 0.4914538 | 0.34751030 | 3.49% |
| Cusp 1-7 | 0.4110000 | 6.45% | 0.4685574 | 0.3313201 | 5.04% | 0.5898000 | 8.05% | 0.7002118 | 0.4951245 | 6.35% | 0.3356667 | 4.23% | 0.3529764 | 0.24959200 | 3.13% |
| Cusp 2-3 | 0.3877333 | 5.93% | 0.4759711 | 0.3365624 | 5.07% | 0.4265333 | 5.67% | 0.5687228 | 0.4021477 | 4.79% | 0.4248667 | 5.11% | 0.5209740 | 0.36838430 | 4.39% |
| Cusp 2-4 | 0.4115333 | 7.48% | 0.5442299 | 0.3848287 | 6.64% | 0.3721333 | 5.80% | 0.4265108 | 0.3015887 | 4.53% | 0.3649333 | 5.74% | 0.4144325 | 0.29304800 | 4.40% |
| Cusp 2-5 | 0.3413333 | 3.92% | 0.4227869 | 0.2989555 | 3.35% | 0.4590769 | 5.00% | 0.5639615 | 0.3987810 | 4.06% | 0.3998333 | 4.16% | 0.5031764 | 0.35579950 | 3.55% |
| Cusp 2-6 | 0.3185000 | 3.86% | 0.3818832 | 0.2700322 | 3.25% | 0.5013333 | 6.15% | 0.599069 | 0.4236058 | 4.66% | 0.2834000 | 2.91% | 0.2931822 | 0.20731110 | 2.14% |
| Cusp 2-7 | 0.1305000 | 3.98% | 0.1305086 | 0.09228353 | 2.80% | 0.2572000 | 7.54% | 0.3247331 | 0.2296210 | 6.08% | 0.2986667 | 8.06% | 0.3016930 | 0.21332920 | 5.68% |
| Cusp 3-4 | 0.3932000 | 6.09% | 0.4637065 | 0.3278900 | 4.95% | 0.3576667 | 4.95% | 0.4365338 | 0.3086760 | 4.20% | 0.4442000 | 6.21% | 0.5531816 | 0.39115850 | 5.31% |
| Cusp 3-5 | 0.3590000 | 9.07% | 0.4887228 | 0.3455792 | 8.40% | 0.2744615 | 8.21% | 0.3551234 | 0.2511102 | 6.44% | 0.2515833 | 6.54% | 0.3105263 | 0.21957530 | 5.71% |
| Cusp 3-6 | 0.3232500 | 5.05% | 0.3642688 | 0.2575769 | 3.94% | 0.3406667 | 6.22% | 0.3811137 | 0.2694881 | 4.56% | 0.2982000 | 5.13% | 0.3402796 | 0.24061400 | 3.99% |
| Cusp 3-7 | 0.2415000 | 3.86% | 0.2809172 | 0.1986385 | 3.09% | 0.4918000 | 6.73% | 0.6245176 | 0.4416006 | 5.67% | 0.3076667 | 3.86% | 0.3628760 | 0.25659210 | 3.26% |
| Cusp 4-5 | 0.2422667 | 4.54% | 0.3481594 | 0.2461858 | 4.37% | 0.2517692 | 4.93% | 0.3736176 | 0.2641875 | 4.52% | 0.3135833 | 5.60% | 0.4158200 | 0.29402910 | 4.70% |
| Cusp 4-6 | 0.1821250 | 5.24% | 0.2033350 | 0.1437796 | 4.31% | 0.2685000 | 8.77% | 0.3381319 | 0.2390953 | 6.46% | 0.1064000 | 2.41% | 0.1363378 | 0.09640539 | 2.01% |
| Cusp 4-7 | 0.3410000 | 13.74% | 0.3628457 | 0.2565707 | 9.96% | 0.2040000 | 7.15% | 0.2443907 | 0.1728103 | 5.47% | 0.1403333 | 3.70% | 0.1623874 | 0.11482520 | 3.04% |
| Cusp 5-6 | 0.1631250 | 4.24% | 0.1832345 | 0.1295663 | 3.41% | 0.1255000 | 4.06% | 0.1647346 | 0.116485 | 3.74% | 0.1910000 | 6.74% | 0.2159903 | 0.15272820 | 4.89% |
| Cusp 5-7 | 0.2735000 | 3.90% | 0.2735041 | 0.1933966 | 2.70% | 0.3704 | 4.64% | 0.4683396 | 0.3311661 | 4.10% | 0.3773333 | 4.53% | 0.4206186 | 0.2974223 | 3.57% |
| Cusp 6-7 | 0.1170000 | 2.37% | 0.1170000 | 0.08273149 | 1.66% | 0.2153333 | 3.44% | 0.2851888 | 0.201659 | 3.23% | 0.05 | 0.69% | 0.05 | 0.03535534 | 0.48% |

|  | | Metameric Comparison | | | | | Antimeric Comparison | | | | |
| --- | --- | --- | --- | --- | --- | --- | --- | --- | --- | --- | --- |
|  |  | dm_2_ Mean | M_1_ Mean | rTEM (%) | V^2^ | p-value^3^ | LM_1_ Mean | RM_1_ Mean | rTEM (%) | V | p-value |
| Trace Area (mm^2^) | Base Crown Area | **73.04** | **93.53** | **19.25** | **6** | **0.00** | **93.86** | **95.78** | **5.87** | **2161** | **0.01** |
|  | Square Root Crown Area | **8.53** | **9.66** | **9.64** | **6** | **0.00** | **9.67** | **9.77** | **3.02** | **2169** | **0.01** |
| Absolute Intercusp Distance (mm) | Cusp 1-2 | **4.51** | **5.95** | **21.58** | **13** | **0.00** | **5.93** | **6.21** | **7.80** | **1485** | **0.00** |
|  | Cusp 1-3 | **4.32** | **5.33** | **19.35** | **50** | **0.00** | 5.26 | 5.23 | 9.25 | 3076.5 | 0.81 |
|  | Cusp 1-4 | **8.10** | **9.35** | **12.17** | **36** | **0.00** | **9.42** | **9.59** | **4.84** | **2111** | **0.01** |
|  | Cusp 1-5 | **8.07** | **8.67** | **10.14** | **185** | **0.00** | 8.79 | 8.81 | 5.06 | 2468 | 0.60 |
|  | Cusp 1-6 | **9.37** | **9.93** | **8.98** | **39** | **0.04** | 10.18 | 10.13 | 3.88 | 675.5 | 0.72 |
|  | Cusp 1-7 | 5.87 | 7.94 | 23.34 | 0 | 0.25 | 8.30 | 8.35 | 4.07 | 72 | 0.57 |
|  | Cusp 2-3 | **6.68** | **8.61** | **20.39** | **9** | **0.00** | **8.54** | **8.69** | **5.84** | **2306** | **0.04** |
|  | Cusp 2-4 | **5.85** | **6.78** | **12.94** | **31** | **0.00** | **6.85** | **6.97** | **6.10** | **2245.5** | **0.02** |
|  | Cusp 2-5 | **8.94** | **10.06** | **11.06** | **56** | **0.00** | **10.23** | **10.42** | **4.21** | **1585.5** | **0.00** |
|  | Cusp 2-6 | **8.38** | **9.65** | **12.21** | **6** | **0.00** | 9.66 | 9.77 | 4.37 | 516 | 0.24 |
|  | Cusp 2-7 | 2.73 | 3.71 | 34.65 | 0 | 0.25 | 4.19 | 3.97 | 8.94 | 126 | 0.08 |
|  | Cusp 3-4 | **6.56** | **7.51** | **12.27** | **49** | **0.00** | 7.55 | 7.60 | 5.53 | 2662.5 | 0.31 |
|  | Cusp 3-5 | 4.00 | 3.84 | 14.51 | 555 | 0.11 | 3.99 | 3.99 | 10.62 | 2574.5 | 0.86 |
|  | Cusp 3-6 | 6.34 | 6.05 | 10.78 | 116 | 0.20 | **6.49** | **6.34** | **5.64** | **857** | **0.03** |
|  | Cusp 3-7 | 6.19 | 7.77 | 18.125 | 0 | 0.25 | 8.16 | 8.10 | 7.92 | 77 | 0.73 |
|  | Cusp 4-5 | **5.69** | **6.02** | **12.34** | **269** | **0.04** | 6.20 | 6.27 | 6.79 | 2078.5 | 0.07 |
|  | Cusp 4-6 | **3.25** | **4.12** | **21.43** | **10** | **0.00** | **3.87** | **4.02** | **10.13** | **414.5** | **0.03** |
|  | Cusp 4-7 | 2.98 | 3.23 | 35.93 | 3 | 1 | **3.44** | **3.86** | **14.36** | **26** | **0.01** |
|  | Cusp 5-6 | **3.78** | **3.23** | **19.25** | **143** | **0.01** | 3.53 | 3.44 | 9.92 | 756 | 0.25 |
|  | Cusp 5-7 | 7.34 | 7.98 | 9.05 | 0 | 0.25 | 8.31 | 8.35 | 4.96 | 67 | 0.68 |
|  | Cusp 6-7 | 6.77 | 6.43 | 3.60 | 1 | 1 | 6.39 | 7.22 | 10.56 | 0 | 0.13 |
| Relative Intercusp Distance^1^  (mm) | Cusp 1-2 | **0.53** | **0.62** | **13.91** | **45** | **0.00** | **0.61** | **0.64** | **6.52** | **1579** | **0.00** |
|  | Cusp 1-3 | **0.51** | **0.55** | **12.63** | **195** | **0.00** | 0.54 | 0.54 | 8.78 | 3488 | 0.14 |
|  | Cusp 1-4 | **0.95** | **0.97** | **5.57** | **250** | **0.01** | 0.97 | 0.98 | 3.31 | 2547 | 0.17 |
|  | Cusp 1-5 | **0.94** | **0.90** | **8.05** | **627** | **0.01** | 0.91 | 0.90 | 4.37 | 3095 | 0.12 |
|  | Cusp 1-6 | **1.10** | **1.01** | **8.84** | **150** | **0.00** | 1.04 | 1.03 | 2.86 | 840 | 0.05 |
|  | Cusp 1-7 | 0.66 | 0.81 | 17.41 | 0 | 0.25 | 0.84 | 0.84 | 3.85 | 75 | 0.67 |
|  | Cusp 2-3 | **0.78** | **0.89** | **12.25** | **61** | **0.00** | 0.88 | 0.89 | 4.57 | 2686 | 0.35 |
|  | Cusp 2-4 | **0.68** | **0.70** | **6.55** | **310** | **0.05** | 0.71 | 0.71 | 5.30 | 2516 | 0.15 |
|  | Cusp 2-5 | 1.04 | 1.04 | 4.59 | 462 | 0.69 | **1.05** | **1.06** | **3.01** | **1899** | **0.02** |
|  | Cusp 2-6 | 0.98 | 0.98 | 5.63 | 72 | 0.58 | 0.99 | 0.99 | 3.22 | 561 | 0.46 |
|  | Cusp 2-7 | 0.31 | 0.38 | 30.91 | 2 | 0.75 | 0.42 | 0.40 | 9.19 | 129 | 0.06 |
|  | Cusp 3-4 | 0.77 | 0.78 | 5.61 | 364 | 0.19 | 0.78 | 0.78 | 4.21 | 3226 | 0.49 |
|  | Cusp 3-5 | **0.47** | **0.40** | **17.76** | **760** | **0.00** | 0.41 | 0.41 | 10.27 | 2863 | 0.43 |
|  | Cusp 3-6 | **0.74** | **0.62** | **15.58** | **168** | **0.00** | **0.66** | **0.62** | **14.81** | **1046** | **0.00** |
|  | Cusp 3-7 | 0.69 | 0.79 | 10.85 | 0 | 0.25 | 0.82 | 0.81 | 7.75 | 80 | 0.83 |
|  | Cusp 4-5 | **0.66** | **0.62** | **9.95** | **648** | **0.00** | 0.64 | 0.64 | 5.97 | 2578 | 0.87 |
|  | Cusp 4-6 | **0.38** | **0.42** | **15.14** | **39** | **0.04** | 0.40 | 0.39 | 16.17 | 544 | 0.19 |
|  | Cusp 4-7 | 0.33 | 0.33 | 36.00 | 3 | 1 | **0.35** | **0.39** | **14.37** | **31** | **0.02** |
|  | Cusp 5-6 | **0.44** | **0.33** | **24.32** | **168** | **0.00** | 0.36 | 0.33 | 17.81 | 863 | 0.06 |
|  | Cusp 5-7 | 0.82 | 0.81 | 5.70 | 4 | 0.75 | 0.84 | 0.84 | 4.19 | 83 | 0.78 |
|  | Cusp 6-7 | 0.76 | 0.64 | 11.42 | 1 | 1 | 0.65 | 0.72 | 8.58 | 0 | 0.13 |

**S4 Table. Results of metameric and antimeric comparative analyses for trace crown area, AICD, and RICD**

^1^Intercusp distance scaled by traced crown base area

^2^V: Paired-samples Wilcoxon signed-rank test statistic

^3^α≤0.05; significant results in bold

|  | | Metameric Comparison | | | | | Antimeric Comparison | | | | |
| --- | --- | --- | --- | --- | --- | --- | --- | --- | --- | --- | --- |
|  |  | dm_2_ Mean | M_1_ Mean | rTEM (%) | V^8^ | p-value^9^ | LM_1_ Mean | RM_1_ Mean | rTEM (%) | V | p-value |
| Absolute Intercusp Area (mm^2^) | Cusps 1-4^5^ | **26.22** | **40.37** | **32.74** | **8** | **0.00** | **40.15** | **41.41** | **9.70** | **2102** | **0.01** |
|  | Cusps 1-5^6^ | **37.23** | **51.57** | **25.48** | **14** | **0.00** | **52.23** | **53.80** | **8.67** | **1812** | **0.01** |
|  | Cusps 1-6^7^ | **43.67** | **57.72** | **24.81** | **0** | **0.00** | 57.85 | 59.72 | 6.98 | 529 | 0.21 |
| Relative Intercusp Area^1^ (mm^2^) | Cusps 1-4 | **2.79** | **3.84** | **24.47** | **11** | **0.00** | **3.80** | **3.88** | **8.21** | **2291** | **0.03** |
|  | Cusps 1-5 | **3.97** | **4.66** | **22.32** | **106** | **0.00** | **4.93** | **5.01** | **7.18** | **1989** | **0.03** |
|  | Cusps 1-6 | **4.61** | **5.40** | **15.55** | **6** | **0.00** | 5.46 | 5.56 | 5.69 | 564 | 0.36 |
| Relative Intercusp Area^2^ (mm^2^) | Cusps 1-4 | **3.07** | **4.17** | **23.73** | **14** | **0.00** | **4.14** | **4.23** | **8.11** | **2251** | **0.02** |
|  | Cusps 1-5 | **4.35** | **5.31** | **16.21** | **25** | **0.00** | **5.37** | **5.47** | **7.07** | **1946** | **0.02** |
|  | Cusps 1-6 | **5.06** | **5.90** | **15.31** | **6** | **0.00** | 5.96 | 6.07 | 5.50 | 559 | 0.33 |
| Relative Intercusp Area^3^ (mm^2^) | Cusps 1-4 | **8.03** | **11.99** | **30.20** | **7** | **0.00** | **11.87** | **12.19** | **8.91** | **2185** | **0.01** |
|  | Cusps 1-5 | **11.40** | **15.29** | **22.69** | **14** | **0.00** | **15.41** | **15.78** | **7.90** | **1896** | **0.01** |
|  | Cusps 1-6 | **13.28** | **17.03** | **21.70** | **0** | **0.00** | 17.06 | 17.53 | 6.25 | 547 | 0.28 |
| Relative Intercusp Area^4^ (mm^2^) | Cusps 1-4 | **9.12** | **12.93** | **26.95** | **14** | **0.00** | **12.85** | **13.18** | **8.93** | **2210** | **0.02** |
|  | Cusps 1-5 | **12.94** | **16.48** | **19.60** | **23** | **0.00** | **16.69** | **17.08** | **7.87** | **1874** | **0.01** |
|  | Cusps 1-6 | **15.14** | **18.31** | **18.63** | **3** | **0.00** | **16.94** | **18.92** | **10.26** | **85** | **0.00** |

**S5. Results of metameric and antimeric comparative analyses for intercusp area**

^1^Intercusp area scaled by BLxMD crown base area

^2^Intercusp area scaled by trace crown base

^3^Intercusp area scaled by BL dimension

^4^Intercusp area scaled by MD dimension

^5^Intercusp area formed by protoconid-metaconid-hypoconid-entoconid polygon

^6^Intercusp area formed by protoconid-metaconid-hypoconid-entoconid-hypoconulid polygon

^7^Intercusp area formed by protoconid-metaconid-hypoconid-entoconid-hypoconulid-cusp6 polygon

^8^V: Paired-samples Wilcoxon signed-rank test statistic

^9^α≤0.05; significant results in bold

**S6 Table. Results of proportional odds regression using tooth dimension and tooth size as the predictor of accessory cusp expression.** Cells highlighted in blue represent dimensions/areas that become significantly longer/larger with increasing accessory cusp expression. Cells highlighted in red represent dimensions/areas that become significantly shorter/smaller with increasing accessory cusp expression.

|  | | dm_2_ | | | LM_1_ | | | RM_1_ | | |
| --- | --- | --- | --- | --- | --- | --- | --- | --- | --- | --- |
|  |  | Cusp 5 | Cusp 6 | Cusp 7 | Cusp 5 | Cusp 6 | Cusp 7 | Cusp 5 | Cusp 6 | Cusp 7 |
| MD | Coefficient | 0.24 | 0.43 | 0.18 | **0.64** | 0.18 | 0.39* | **1.07** | 0.30 | 0.38 |
|  | LRχ^1^ | 0.26 | 0.89 | 0.04 | **6.84** | 0.59 | 1.27 | **15.56** | 1.34 | 1.13 |
|  | p-value^2^ | 0.61 | 0.34 | 0.85 | **0.01** | 0.44 | 0.26 | **0.00** | 0.25 | 0.29 |
|  | Odds ratio^3^ | - | - | - | **1.07** | - | - | **1.11** | - | - |
| BL | Coefficient | -0.05 | 0.38 | 2.70 | 0.42* | **0.66** | 0.41 | 0.33 | **0.79** | **0.83** |
|  | LRχ | 0.01 | 0.51 | 3.77 | 2.08 | **5.24** | 1.14 | 1.25 | **6.53** | **4.12** |
|  | p-value | 0.93 | 0.47 | 0.05 | 0.15 | **0.02** | 0.29 | 0.26 | **0.01** | **0.04** |
|  | Odds ratio | - | - | - | - | **1.07** | - | - | **1.08** | **1.09** |
| BLxMD | Coefficient | 0.01 | 0.03 | 0.13 | **0.03** | 0.02 | 0.02 | **0.04** | **0.03** | 0.03 |
|  | LRχ | 0.05 | 1.06 | 2.00 | **4.68** | 2.72 | 1.42 | **7.16** | **3.89** | 2.75 |
|  | p-value | 0.82 | 0.30 | 0.16 | **0.03** | 0.10 | 0.23 | **0.01** | **0.05** | 0.10 |
|  | Odds ratio | - | - | - | **1.00** | - | - | **1.00** | **1.00** | - |
| Trace | Coefficient | -0.00 | 0.03 | 0.21 | 0.03 | 0.02 | 0.04 | **0.04** | 0.03 | 0.04 |
|  | LRχ | 0.01 | 0.61 | 3.34 | 2.95 | 1.17 | 2.69 | **4.87** | 2.61 | 3.20 |
|  | p-value | 0.91 | 0.43 | 0.07 | 0.09 | 0.28 | 0.10 | **0.03** | 0.11 | 0.07 |
|  | Odds ratio | - | - | - | - | - | - | **1.00** | - | - |

^*^Values violating the assumption of proportionality are marked with an asterisk, these results should be interpreted with caution

^1^LRχ: Likelihood ratio chi-square test statistic

^2^α≤0.05; significant results in bold

^3^Odds ratio obtained by dividing the model coefficient by 10 to scale the odds ratio to a 0.10-unit change and exponentiating the quotient

**S7 Table. Results of proportional odds regression when using tooth dimension and size as the predictor of accessory cusp presence (dichotomized).** Cells highlighted in blue represent dimensions/areas that become significantly longer/larger with accessory cusp development. Cells highlighted in red represent dimensions/areas that become significantly shorter/smaller with accessory cusp development.

|  | | dm_2_ | | | LM_1_ | | | RM_1_ | | |
| --- | --- | --- | --- | --- | --- | --- | --- | --- | --- | --- |
|  |  | Cusp 5 | Cusp 6 | Cusp 7 | Cusp 5 | Cusp 6 | Cusp 7 | Cusp 5 | Cusp 6 | Cusp 7 |
| MD | Coefficient | - | 0.71 | 0.18 | **1.53** | 0.22 | 0.40 | **2.18** | 0.25 | 0.38 |
|  | p-value^1^ | - | 0.17 | 0.86 | **0.01** | 0.41 | 0.26 | **0.00** | 0.38 | 0.29 |
|  | Odds ratio^2^ | - | - | - | **1.17** | - | - | **1.24** | - | - |
| BL | Coefficient | - | 0.38 | 2.82 | **1.98** | 0.57 | 0.45 | **1.87** | 0.51 | **0.84** |
|  | p-value | - | 0.50 | 0.10 | **0.02** | 0.07 | 0.25 | **0.01** | 0.12 | **0.05** |
|  | Odds ratio | - | - | - | **1.22** | - | - | **1.21** | - | **1.09** |
| BLxMD | Coefficient | - | 0.04 | 0.13 | **0.10** | 0.02 | 0.02 | **0.12** | 0.02 | 0.03 |
|  | p-value | - | 0.21 | 0.22 | **0.01** | 0.15 | 0.21 | **0.00** | 0.19 | 0.10 |
|  | Odds ratio | - | - | - | **1.01** | - | - | **1.01** | - | - |
| Trace | Coefficient | - | 0.03 | 0.21 | 0.08 | 0.02 | 0.04 | **0.11** | 0.02 | 0.04 |
|  | p-value | - | 0.38 | 0.14 | 0.06 | 0.35 | 0.09 | **0.02** | 0.33 | 0.07 |
|  | Odds ratio | - | - | - | - | - | - | **1.01** | - | - |

^1^α≤0.05; significant results in bold

^2^Odds ratio obtained by dividing the model coefficient by 10 to scale the odds ratio to a 0.10-unit change and exponentiating the quotient

|  | | | Cusp 1-2 | Cusp 1-3 | Cusp 1-4 | Cusp 1-5 | Cusp 1-6 | Cusp 2-3 | Cusp 2-4 | Cusp 2-5 | Cusp 2-6 | Cusp 3-4 | Cusp 3-5 | Cusp 3-6 | Cusp 4-5 | Cusp 4-6 | Cusp 5-6 |
| --- | --- | --- | --- | --- | --- | --- | --- | --- | --- | --- | --- | --- | --- | --- | --- | --- | --- |
| dm_2_ | Cusp 5 | Coefficient | 0.29 | -0.30* | 0.30* | - | - | -0.05 | 0.34 | - | - | 0.14 | - | - | - | - | - |
|  |  | LRχ^1^ | 0.41 | 0.43 | 0.76 | - | - | 0.02 | 0.81 | - | - | 0.11 | - | - | - | - | - |
|  |  | p-value^2^ | 0.52 | 0.51 | 0.38 | - | - | 0.89 | 0.37 | - | - | 0.75 | - | - | - | - | - |
|  |  | Odds ratio^3^ | - | - | - | - | - | - | - | - | - | - | - | - | - | - | - |
|  | Cusp 6 | Coefficient | -0.19 | 0.35 | -0.03 | 0.16 | - | 0.19 | -0.28 | 0.36 | - | 0.56 | -0.52 | - | **1.75** | - | - |
|  |  | LRχ | 0.14 | 0.63 | 0.01 | 0.18 | - | 0.33 | 0.55 | 1.00 | - | 1.48 | 0.59 | - | **11.57** | - | - |
|  |  | p-value | 0.70 | 0.43 | 0.93 | 0.67 | - | 0.57 | 0.46 | 0.32 | - | 0.22 | 0.44 | - | **0.00** | - | - |
|  |  | Odds ratio | - | - | - | - | - | - | - | - | - | - | - | - | **1.19** | - | - |
|  | Cusp 7 | Coefficient | -0.20 | -0.10 | -0.11 | -0.42 | -0.55 | 0.04 | 0.21 | 0.17 | -0.19 | -0.17 | -1.75 | -0.21 | 0.55 | -2.29 | 0.53 |
|  |  | LRχ | 0.05 | 0.02 | 0.02 | 0.25 | 0.39 | 0.00 | 0.09 | 0.06 | 0.04 | 0.04 | 1.20 | 0.02 | 0.40 | 2.34 | 0.18 |
|  |  | p-value | 0.82 | 0.90 | 0.88 | 0.62 | 0.53 | 0.95 | 0.76 | 0.81 | 0.84 | 0.85 | 0.27 | 0.88 | 0.53 | 0.13 | 0.67 |
|  |  | Odds ratio | - | - | - | - | - | - | - | - | - | - | - | - | - | - | - |
| LM_1_ | Cusp 5 | Coefficient | 0.13 | **-0.65*** | 0.21 | - | - | -0.28 | 0.07 | - | - | 0.33 | - | - | - | - | - |
|  |  | LRχ | 0.28 | **6.85** | 1.26 | - | - | 2.05 | 0.13 | - | - | 2.19 | - | - | - | - | - |
|  |  | p-value | 0.59 | **0.01** | 0.26 | - | - | 0.15 | 0.72 | - | - | 0.14 | - | - | - | - | - |
|  |  | Odds ratio | - | **0.94** | - | - | - | - | - | - | - | - | - | - | - | - | - |
|  | Cusp 6 | Coefficient | -0.1 | -0.14 | -0.19 | -0.21 | **-** | 0.18 | -0.38 | 0.21 | **-** | **0.55** | -0.47 | **-** | **1.24** | - | - |
|  |  | LRχ | 0.00 | 0.37 | 1.01 | 1.00 | **-** | 0.85 | 3.69 | 1.11 | **-** | **5.94** | 2.72 | **-** | **24.07** | - | - |
|  |  | p-value | 0.97 | 0.55 | 0.32 | 0.32 | **-** | 0.36 | 0.05 | 0.29 | **-** | **0.01** | 0.10 | **-** | **0.00** | - | - |
|  |  | Odds ratio | - | - | - | - | - | - | - | - | - | **1.06** | - | - | **1.13** | - | - |
|  | Cusp 7 | Coefficient | -0.10 | -0.28 | 0.10 | -0.08 | -0.70* | -0.27 | 0.41 | -0.08 | -0.55 | -0.24 | -0.05 | **-1.21** | -0.36 | -0.51 | -0.12 |
|  |  | LRχ | 0.08 | 0.75 | 0.15 | 0.07 | 2.49 | 1.06 | 2.27 | 0.09 | 2.07 | 0.69 | 0.02 | **4.00** | 1.75 | 0.92 | 0.04 |
|  |  | p-value | 0.78 | 0.39 | 0.70 | 0.79 | 0.11 | 0.30 | 0.13 | 0.76 | 0.15 | 0.41 | 0.90 | **0.05** | 0.19 | 0.34 | 0.83 |
|  |  | Odds ratio | - | - | - | - | - | - | - | - | - | - | - | **0.89** | - | - | - |
| RM_1_ | Cusp 5 | Coefficient | 0.42 | **-0.89*** | 0.35 | - | - | -0.09 | 0.40 | - | - | **0.48*** | - | - | - | - | - |
|  |  | LRχ | 2.02 | **9.73** | 3.06 | - | - | 0.16 | 3.75 | - | - | **4.08** | - | - | - | - | - |
|  |  | p-value | 0.16 | **0.00** | 0.08 | - | - | 0.69 | 0.05 | - | - | **0.04** | - | - | - | - | - |
|  |  | Odds ratio | - | **0.91** | - | - | - | - | - | - | - | **1.05** | - | - | - | - | - |
|  | Cusp 6 | Coefficient | -0.19 | -0.12 | -0.17 | -0.25 | - | 0.33 | **-0.47** | 0.30 | - | **0.86** | **-0.73** | - | **1.70** | - | - |
|  |  | LRχ | 0.38 | 0.19 | 0.64 | 1.11 | - | 1.73 | **4.48** | 1.63 | - | **11.56** | **5.29** | - | **38.17** | - | - |
|  |  | p-value | 0.54 | 0.67 | 0.43 | 0.29 | - | 0.19 | **0.03** | 0.20 | - | **0.00** | **0.02** | - | **0.00** | - | - |
|  |  | Odds ratio | - | - | - | - | - | - | **0.95** | - | - | **1.09** | **0.93** | - | **1.19** | - | - |
|  | Cusp 7 | Coefficient | 0.11 | -0.19 | 0.35 | 0.25 | 0.27 | -0.17 | **0.80** | -0.00 | 0.32 | -0.22 | 0.59 | -0.00 | **-0.88** | -0.49 | -0.18* |
|  |  | LRχ | 0.08 | 0.25 | 1.45 | 0.66 | 0.17 | 0.26 | **6.30** | 0.00 | 0.25 | 0.46 | 2.12 | 0.00 | **7.03** | 0.80 | 0.08 |
|  |  | p-value | 0.78 | 0.62 | 0.23 | 0.42 | 0.68 | 0.61 | **0.01** | 1 | 0.62 | 0.50 | 0.15 | 1 | **0.01** | 0.37 | 0.78 |
|  |  | Odds ratio | - | - | - | - | - | - | **1.08** | - | - | - | - | - | **0.92** | - | - |

**S8 Table. Results of proportional ordinal regression when using AICD as a predictor of lower molar accessory cusp trait expression.** Cells highlighted in blue represent intercusp distances that become significantly longer with increasing accessory cusp expression. Cells highlighted in red represent intercusp distances that become significantly shorter with increasing accessory cusp expression.

^*^Values violating the assumption of proportionality are marked with an asterisk, these results should be interpreted with caution

^1^LRχ: Likelihood ratio chi-square test statistic

^2^α≤0.05; significant results in bold

^3^Odds ratio obtained by dividing the model coefficient by 10 to scale the odds ratio to a 0.10-unit change and exponentiating the quotient

|  | | | Cusp 1-2 | Cusp 1-3 | Cusp 1-4 | Cusp 1-5 | Cusp 1-6 | Cusp 2-3 | Cusp 2-4 | Cusp 2-5 | Cusp 2-6 | Cusp 3-4 | Cusp 3-5 | Cusp 3-6 | Cusp 4-5 | Cusp 4-6 | Cusp 5-6 |
| --- | --- | --- | --- | --- | --- | --- | --- | --- | --- | --- | --- | --- | --- | --- | --- | --- | --- |
| dm_2_ | Cusp 5 | Coefficient | 3.68 | -2.92* | 3.77* | - | - | -0.45 | 3.57 | - | - | 2.19 | - | - | - | - | - |
|  |  | LRχ^1^ | 0.67 | 0.47 | 1.25 | - | - | 0.01 | 1.11 | - | - | 0.23 | - | - | - | - | - |
|  |  | p-value^2^ | 0.41 | 0.49 | 0.26 | - | - | 0.90 | 0.29 | - | - | 0.63 | - | - | - | - | - |
|  |  | Odds ratio^3^ | - | - | - | - | - | - | - | - | - | - | - | - | - | - | - |
|  | Cusp 6 | Coefficient | -3.70 | 2.17 | -2.08 | -0.00* | - | 0.96 | -4.30 | 3.26* | - | 4.45 | -7.85* | - | **18.99*** | - | - |
|  |  | LRχ | 0.60 | 0.28 | 0.37 | 0.00 | - | 0.07 | 1.47 | 0.59 | - | 0.85 | 1.46 | - | **13.13** | - | - |
|  |  | p-value | 0.44 | 0.60 | 0.54 | 1 | - | 0.79 | 0.23 | 0.44 | - | 0.36 | 0.23 | - | **0.00** | - | - |
|  |  | Odds ratio | - | - | - | - | - | - | - | - | - | - | - | - | **6.68** | - | - |
|  | Cusp 7 | Coefficient | -8.72 | -5.19 | -7.27 | -16.76 | -14.56 | -4.09 | -1.63 | -6.72 | -13.66 | -10.00* | **-32.19** | -12.58 | -0.43 | -28.48 | 1.09 |
|  |  | LRχ | 0.78 | 0.40 | 1.04 | 2.91 | 2.18 | 0.35 | 0.07 | 0.67 | 1.15 | 1.22 | **4.14** | 0.77 | 0.00 | 3.58 | 0.01 |
|  |  | p-value | 0.38 | 0.53 | 0.31 | 0.09 | 0.14 | 0.55 | 0.79 | 0.41 | 0.28 | 0.27 | **0.04** | 0.38 | 0.95 | 0.06 | 0.92 |
|  |  | Odds ratio | - | - | - | - | - | - | - | - | - | - | **0.04** | - | - | - | - |
| LM_1_ | Cusp 5 | Coefficient | -1.19 | **-9.52*** | 0.23* | - | - | **-7.27** | -0.90 | - | - | 2.20 | - | - | - | - | - |
|  |  | LRχ | 0.18 | **13.79** | 0.01 | - | - | **9.19** | 0.20 | - | - | 0.63 | - | - | - | - | - |
|  |  | p-value | 0.68 | **0.00** | 0.92 | - | - | **0.00** | 0.66 | - | - | 0.43 | - | - | - | - | - |
|  |  | Odds ratio | - | **0.39** | - | - | - | **0.48** | - | - | - | - | - | - | - | - | - |
|  | Cusp 6 | Coefficient | -2.06 | -2.59 | **-5.40** | -4.32 | **-** | 1.07 | **-5.78** | 2.35 | **-** | **8.02** | **-6.04** | **-** | **17.21** | - | - |
|  |  | LRχ | 0.46 | 1.17 | **4.48** | 2.88 | **-** | 0.20 | **7.00** | 0.78 | **-** | **7.09** | **4.05** | **-** | **31.32** | - | - |
|  |  | p-value | 0.50 | 0.28 | **0.03** | 0.09 | **-** | 0.66 | **0.01** | 0.38 | **-** | **0.01** | **0.04** | **-** | **0.00** | - | - |
|  |  | Odds ratio | - | - | **0.58** | - | - | - | **0.56** | - | - | **2.23** | **0.55** | - | **5.59** | - | - |
|  | Cusp 7 | Coefficient | -4.97 | -5.41 | -2.04 | -4.52 | **-13.11** | **-6.70** | 2.40 | -5.37 | -7.35 | **-7.70*** | -2.54 | **-13.92** | **-5.83*** | -5.44 | 0.39 |
|  |  | LRχ | 1.54 | 2.64 | 0.42 | 1.82 | **3.88** | **5.01** | 0.71 | 2.52 | 2.29 | **4.36** | 0.44 | **4.28** | **3.99** | 0.86 | 0.00 |
|  |  | p-value | 0.21 | 0.10 | 0.52 | 0.18 | **0.5** | **0.03** | 0.40 | 0.11 | 0.13 | **0.04** | 0.51 | **0.04** | **0.05** | 0.35 | 0.95 |
|  |  | Odds ratio | - | - | - | - | **0.27** | **0.51** | - | - | - | **0.46** | - | **0.25** | **0.56** | - | - |
| RM_1_ | Cusp 5 | Coefficient | 0.23 | **-12.39*** | 0.94 | - | - | **-7.43** | 1.93 | - | - | 3.67 | - | - | - | - | - |
|  |  | LRχ | 0.00 | **19.32** | 0.15 | - | - | **6.06** | 0.84 | - | - | 1.30 | - | - | - | - | - |
|  |  | p-value | 0.95 | **0.00** | 0.70 | - | - | **0.01** | 0.36 | - | - | 0.25 | - | - | - | - | - |
|  |  | Odds ratio | - | **0.29** | - | - | - | **0.48** | - | - | - | - | - | - | - | - | - |
|  | Cusp 6 | Coefficient | -6.28 | -3.10 | **-5.64** | **-5.62** | - | 1.08 | **-6.92** | 2.33* | - | **12.72** | **-10.26** | - | **20.54** | - | - |
|  |  | LRχ | 3.15 | 1.34 | **4.54** | **3.95** | - | 0.13 | **8.81** | 0.59 | - | **13.22** | **8.56** | - | **42.68** | - | - |
|  |  | p-value | 0.08 | 0.25 | **0.03** | **0.05** | - | 0.72 | **0.00** | 0.44 | - | **0.00** | **0.00** | - | **0.00** | - | - |
|  |  | Odds ratio | - | - | **0.57** | **0.57** | - | - | **0.50** | - | - | **3.57** | **0.36** | - | **7.80** | - | - |
|  | Cusp 7 | Coefficient | -3.87 | -5.10 | 0.27 | -0.53 | -4.57 | **-8.31** | 5.53 | -5.26 | -3.02 | **-10.62** | 4.31 | -2.85 | **-15.00** | -5.12 | -4.60 |
|  |  | LRχ | 0.73 | 1.78 | 0.01 | 0.02 | 0.37 | **4.30** | 2.92 | 1.90 | 0.18 | **5.51** | 0.95 | 1.71 | **14.75** | 2.82 | 1.60 |
|  |  | p-value | 0.39 | 0.18 | 0.93 | 0.88 | 0.54 | **0.04** | 0.09 | 0.17 | 0.68 | **0.02** | 0.33 | 0.19 | **0.00** | 0.09 | 0.21 |
|  |  | Odds ratio | - | - | - | - | - | **0.44** | - | - | - | **0.35** | - | - | **0.22** | - | - |

**S9 Table. Results of proportional ordinal regression when using RICD scaled by trace crown area as a predictor of lower molar accessory cusp trait expression.** Cells highlighted in blue represent intercusp distances that become significantly longer with increasing accessory cusp expression. Cells highlighted in red represent intercusp distances that become significantly shorter with increasing accessory cusp expression.

^*^Values violating the assumption of proportionality are marked with an asterisk, these results should be interpreted with caution

^1^LRχ: Likelihood ratio chi-square test statistic

^2^α≤0.05; significant results in bold

^3^Odds ratio obtained by dividing the model coefficient by 10 to scale the odds ratio to a 0.10-unit change and exponentiating the quotient

|  | | | Cusp 1-2 | Cusp 1-3 | Cusp 1-4 | Cusp 1-5 | Cusp 1-6 | Cusp 2-3 | Cusp 2-4 | Cusp 2-5 | Cusp 2-6 | Cusp 3-4 | Cusp 3-5 | Cusp 3-6 | Cusp 4-5 | Cusp 4-6 | Cusp 5-6 |
| --- | --- | --- | --- | --- | --- | --- | --- | --- | --- | --- | --- | --- | --- | --- | --- | --- | --- |
| dm_2_ | Cusp 5 | Coefficient | 1.06 | -1.39* | 1.30* | - | - | -0.38 | 1.19 | - | - | 0.40 | - | - | - | - | - |
|  |  | LRχ^1^ | 0.40 | 0.70 | 0.88 | - | - | 0.08 | 0.79 | - | - | 0.05 | - | - | - | - | - |
|  |  | p-value^2^ | 0.52 | 0.40 | 0.35 | - | - | 0.77 | 0.37 | - | - | 0.82 | - | - | - | - | - |
|  |  | Odds ratio^3^ | - | - | - | - | - | - | - | - | - | - | - | - | - | - | - |
|  | Cusp 6 | Coefficient | -1.27 | 1.07 | -0.77 | 0.23 | - | 0.51 | -1.62 | 1.45 | - | 2.00 | -2.78* | - | **7.49** | - | - |
|  |  | LRχ | 0.47 | 0.44 | 0.26 | 0.02 | - | 0.15 | 1.27 | 0.86 | - | 1.20 | 1.28 | - | **13.81** | - | - |
|  |  | p-value | 0.49 | 0.51 | 0.61 | 0.88 | - | 0.70 | 0.26 | 0.35 | - | 0.27 | 0.26 | - | **0.00** | - | - |
|  |  | Odds ratio | - | - | - | - | - | - | - | - | - | - | - | - | **2.11** | - | - |
|  | Cusp 7 | Coefficient | -0.86 | -0.50 | -0.65 | -2.58 | -2.08 | 0.12 | 0.74 | 0.75 | 0.13 | -0.90 | -6.70 | 0.41 | 2.24 | -7.07 | 2.17 |
|  |  | LRχ | 0.08 | 0.03 | 0.06 | 0.51 | 0.30 | 0.00 | 0.09 | 0.06 | 0.00 | 0.07 | 1.46 | 0.01 | 0.48 | 2.08 | 0.30 |
|  |  | p-value | 0.78 | 0.87 | 0.81 | 0.47 | 0.59 | 0.96 | 0.77 | 0.81 | 0.97 | 0.79 | 0.23 | 0.93 | 0.49 | 0.15 | 0.59 |
|  |  | Odds ratio | - | - | - | - | - | - | - | - | - | - | - | - | - | - | - |
| LM_1_ | Cusp 5 | Coefficient | -0.23 | **-3.28*** | 0.22* | - | - | **-2.22** | -0.23 | - | - | 0.76 | - | - | - | - | - |
|  |  | LRχ | 0.06 | **13.06** | 0.08 | - | - | **7.56** | 0.11 | - | - | 0.71 | - | - | - | - | - |
|  |  | p-value | 0.81 | **0.00** | 0.78 | - | - | **0.01** | 0.74 | - | - | 0.40 | - | - | - | - | - |
|  |  | Odds ratio | - | **0.72** | - | - | - | **0.80** | - | - | - | - | - | - | - | - | - |
|  | Cusp 6 | Coefficient | -0.30 | -0.71 | -1.27 | -1.14 | - | 0.65 | **-1.79** | 1.17 | - | **2.61** | -1.90 | - | **5.48** | - | - |
|  |  | LRχ | 0.09 | 0.70 | 2.36 | 1.67 | - | 0.65 | **5.55** | 1.77 | - | **7.66** | 3.26 | - | **30.36** | - | - |
|  |  | p-value | 0.77 | 0.40 | 0.12 | 0.20 | - | 0.42 | **0.02** | 0.18 | - | **0.01** | 0.07 | - | **0.00** | - | - |
|  |  | Odds ratio | - | - | - | - | - | - | **0.84** | - | - | **1.30** | - | - | **1.73** | - | - |
|  | Cusp 7 | Coefficient | -0.91 | -1.42 | 0.05 | -0.80 | -3.80 | -1.66 | 1.38 | -0.90 | -2.55 | -1.63 | -0.41 | **-4.92*** | -1.54 | -1.67 | -0.11 |
|  |  | LRχ | 0.45 | 1.47 | 0.00 | 0.44 | 3.28 | 2.60 | 1.80 | 0.60 | 2.34 | 1.87 | 0.09 | **4.36** | 2.47 | 0.74 | 0.00 |
|  |  | p-value | 0.50 | 0.23 | 0.96 | 0.51 | 0.07 | 0.11 | 0.18 | 0.44 | 0.13 | 0.17 | 0.76 | **0.04** | 0.12 | 0.39 | 0.96 |
|  |  | Odds ratio | - | - | - | - | - | - | - | - | - | - | - | **0.61** | - | - | - |
| RM_1_ | Cusp 5 | Coefficient | 0.22 | **-4.45*** | 0.43 | - | - | **-2.06** | 0.76 | - | - | 1.04* | - | - | - | - | - |
|  |  | LRχ | 0.04 | **19.50** | 0.26 | - | - | **4.59** | 1.04 | - | - | 1.13 | - | - | - | - | - |
|  |  | p-value | 0.85 | **0.00** | 0.61 | - | - | **0.03** | 0.31 | - | - | 0.29 | - | - | - | - | - |
|  |  | Odds ratio | - | **0.64** | - | - | - | **0.81** | - | - | - | - | - | - | - | - | - |
|  | Cusp 6 | Coefficient | -1.32 | -0.69 | -1.26 | -1.41 | - | 1.06 | **-2.10** | 1.67 | - | **4.07** | **-3.18** | - | **7.24** | - | - |
|  |  | LRχ | 1.22 | 0.54 | 1.94 | 1.95 | - | 1.15 | **6.57** | 2.47 | - | **14.67** | **6.83** | - | **46.09** | - | - |
|  |  | p-value | 0.27 | 0.46 | 0.16 | 0.16 | - | 0.28 | **0.01** | 0.12 | - | **0.00** | **0.01** | - | **0.00** | - | - |
|  |  | Odds ratio | - | - | - | - | - | - | **0.81** | - | - | **1.50** | **0.73** | - | **2.06** | - | - |
|  | Cusp 7 | Coefficient | -0.08 | -1.02 | 1.22 | 0.89 | 1.48 | -1.35 | **2.93** | -0.53 | 1.61 | -1.70 | 2.18 | -0.07* | **-3.92** | -1.73 | -0.68* |
|  |  | LRχ | 0.00 | 0.61 | 1.03 | 0.47 | 0.24 | 1.08 | **6.02** | 0.15 | 0.34 | 1.54 | 2.04 | 0.00 | **9.73** | 0.73 | 0.10 |
|  |  | p-value | 0.96 | 0.44 | 0.31 | 0.49 | 0.62 | 0.30 | **0.01** | 0.70 | 0.56 | 0.22 | 0.15 | 0.98 | **0.00** | 0.39 | 0.76 |
|  |  | Odds ratio | - | - | - | - | - | - | **1.34** | - | - | - | - | - | **0.68** | - | - |

**S10 Table. Results of proportional ordinal regression when using RICD scaled by mesiodistal tooth dimension as a predictor of lower molar accessory cusp trait expression.** Cells highlighted in blue represent intercusp distances that become significantly longer with increasing accessory cusp expression. Cells highlighted in red represent intercusp distances that become significantly shorter with increasing accessory cusp expression.

^*^Values violating the assumption of proportionality are marked with an asterisk, these results should be interpreted with caution

^1^LRχ: Likelihood ratio chi-square test statistic

^2^α≤0.05; significant results in bold

^3^Odds ratio obtained by dividing the model coefficient by 10 to scale the odds ratio to a 0.10-unit change and exponentiating the quotient

**S11 Table. Results of proportional ordinal regression when using RICD scaled by buccolingual tooth dimension as a predictor of lower molar accessory cusp trait expression.** Cells highlighted in blue represent intercusp distances that become significantly longer with increasing accessory cusp expression. Cells highlighted in red represent intercusp distances that become significantly shorter with increasing accessory cusp expression.

|  | | | Cusp 1-2 | Cusp 1-3 | Cusp 1-4 | Cusp 1-5 | Cusp 1-6 | Cusp 2-3 | Cusp 2-4 | Cusp 2-5 | Cusp 2-6 | Cusp 3-4 | Cusp 3-5 | Cusp 3-6 | Cusp 4-5 | Cusp 4-6 | Cusp 5-6 |
| --- | --- | --- | --- | --- | --- | --- | --- | --- | --- | --- | --- | --- | --- | --- | --- | --- | --- |
| dm_2_ | Cusp 5 | Coefficient | 0.92 | -0.85* | 0.89* | - | - | -0.16 | 1.00 | - | - | 0.49 | - | - | - | - | - |
|  |  | LRχ^1^ | 0.46 | 0.43 | 0.81 | - | - | 0.02 | 0.87 | - | - | 0.13 | - | - | - | - | - |
|  |  | p-value^2^ | 0.50 | 0.51 | 0.37 | - | - | 0.88 | 0.35 | - | - | 0.72 | - | - | - | - | - |
|  |  | Odds ratio^3^ | - | - | - | - | - | - | - | - | - | - | - | - | - | - | - |
|  | Cusp 6 | Coefficient | -0.82 | 0.78 | -0.34 | 0.21 | - | 0.42 | -1.05 | 0.94 | - | 1.46 | -2.05* | - | **5.63*** | - | - |
|  |  | LRχ | 0.30 | 0.39 | 0.11 | 0.04 | - | 0.17 | 0.89 | 0.69 | - | 1.06 | 1.00 | - | **12.12** | - | - |
|  |  | p-value | 0.58 | 0.53 | 0.74 | 0.85 | - | 0.68 | 0.34 | 0.41 | - | 0.30 | 0.32 | - | **0.00** | - | - |
|  |  | Odds ratio | - | - | - | - | - | - | - | - | - | - | - | - | **1.76** | - | - |
|  | Cusp 7 | Coefficient | -1.76 | -1.07 | -1.40 | -2.98 | -3.28* | -0.73 | -0.16 | -0.93 | -2.51 | -2.26 | -9.19 | -3.81 | 0.41 | -9.68 | 0.42 |
|  |  | LRχ | 0.39 | 0.20 | 0.45 | 1.30 | 1.43 | 0.12 | 0.01 | 0.16 | 0.65 | 0.62 | 3.13 | 0.70 | 0.02 | 3.57 | 0.01 |
|  |  | p-value | 0.53 | 0.66 | 0.50 | 0.26 | 0.23 | 0.73 | 0.93 | 0.69 | 0.42 | 0.43 | 0.08 | 0.40 | 0.88 | 0.06 | 0.91 |
|  |  | Odds ratio | - | - | - | - | - | - | - | - | - | - | - | - | - | - | - |
| LM_1_ | Cusp 5 | Coefficient | 0.08 | **-2.50*** | 0.45 | - | - | **-1.54** | -0.02 | - | - | 1.04 | - | - | - | - | - |
|  |  | LRχ | 0.01 | **9.99** | 0.45 | - | - | **4.85** | 0.00 | - | - | 1.61 | - | - | - | - | - |
|  |  | p-value | 0.92 | **0.00** | 0.5 | - | - | **0.03** | 0.98 | - | - | 0.20 | - | - | - | - | - |
|  |  | Odds ratio | - | **0.78** | - | - | - | **0.86** | - | - | - | - | - | - | - | - | - |
|  | Cusp 6 | Coefficient | -0.67 | -0.86 | **-1.40** | -1.27 | - | 0.18 | **-1.71** | 0.40 | - | **1.87** | **-1.92** | - | **4.54** | - | - |
|  |  | LRχ | 0.56 | 1.34 | **3.99** | 3.01 | - | 0.06 | **6.76** | 0.31 | - | **4.73** | **4.26** | - | **25.51** | - | - |
|  |  | p-value | 0.46 | 0.25 | **0.05** | 0.08 | - | 0.80 | **0.01** | 0.58 | - | **0.03** | **0.04** | - | **0.00** | - | - |
|  |  | Odds ratio | - | - | **0.87** | - | - | - | **0.84** | - | - | **1.21** | **0.83** | - | **1.57** | - | - |
|  | Cusp 7 | Coefficient | -0.77 | -1.18 | 0.06 | -0.57 | -2.61 | -1.35 | 1.10 | -0.68 | -1.83 | -1.42 | -0.32 | -4.00* | -1.50 | -1.54 | -0.05 |
|  |  | LRχ | 0.39 | 1.33 | 0.00 | 0.34 | 2.42 | 2.26 | 1.61 | 0.48 | 1.80 | 1.74 | 0.07 | 3.77 | 2.63 | 0.74 | 0.00 |
|  |  | p-value | 0.53 | 0.25 | 0.95 | 0.56 | 0.12 | 0.13 | 0.20 | 0.49 | 0.18 | 0.19 | 0.79 | 0.05 | 0.10 | 0.39 | 0.98 |
|  |  | Odds ratio | - | - | - | - | - | - | - | - | - | - | - | - | - | - | - |
| RM_1_ | Cusp 5 | Coefficient | 1.08 | **-3.12*** | 0.97 | - | - | -0.85 | 1.03 | - | - | **1.77** | - | - | - | - | - |
|  |  | LRχ | 1.14 | **12.48** | 1.98 | - | - | 1.02 | 2.55 | - | - | **3.97** | - | - | - | - | - |
|  |  | p-value | 0.29 | **0.00** | 0.16 | - | - | 0.31 | 0.11 | - | - | **0.05** | - | - | - | - | - |
|  |  | Odds ratio | - | **0.73** | - | - | - | - | - | - | - | **1.19** | - | - | - | - | - |
|  | Cusp 6 | Coefficient | -1.64 | -0.91 | -1.39 | -1.45 | - | 0.41 | **-1.99** | 0.50* | - | **3.08** | **-2.92** | - | **6.06** | - | - |
|  |  | LRχ | 2.38 | 1.12 | 3.32 | 3.27 | - | 0.22 | **7.73** | 0.36 | - | **10.22** | **7.68** | - | **39.00** | - | - |
|  |  | p-value | 0.12 | 0.29 | 0.07 | 0.07 | - | 0.64 | **0.01** | 0.55 | - | **0.00** | **0.01** | - | **0.00** | - | - |
|  |  | Odds ratio | - | - | - | - | - | - | **0.82** | - | - | **1.36** | **0.75** | - | **1.83** | - | - |
|  | Cusp 7 | Coefficient | -0.54 | -1.19 | 0.52 | 0.22 | -0.24 | -1.58 | 1.91 | -0.83 | 0.04 | -1.90 | 1.51 | -0.64* | **-3.96** | -2.29 | -0.88* |
|  |  | LRχ | 0.16 | 1.03 | 0.28 | 0.05 | 0.01 | 1.91 | 3.74 | 0.61 | 0.00 | 2.41 | 1.29 | 0.09 | **11.50** | 1.38 | 0.20 |
|  |  | p-value | 0.69 | 0.31 | 0.60 | 0.83 | 0.92 | 0.17 | 0.05 | 0.43 | 0.99 | 0.12 | 0.26 | 0.76 | **0.00** | 0.24 | 0.66 |
|  |  | Odds ratio | - | - | - | - | - | - | - | - | - | - | - | - | **0.67** | - | - |

^*^Values violating the assumption of proportionality are marked with an asterisk, these results should be interpreted with caution

^1^LRχ: Likelihood ratio chi-square test statistic

^2^α≤0.05; significant results in bold

^3^Odds ratio obtained by dividing the model coefficient by 10 to scale the odds ratio to a 0.10-unit change and exponentiating the quotient

|  | | | Cusp 1-2 | Cusp 1-3 | Cusp 1-4 | Cusp 1-5 | Cusp 1-6 | Cusp 2-3 | Cusp 2-4 | Cusp 2-5 | Cusp 2-6 | Cusp 3-4 | Cusp 3-5 | Cusp 3-6 | Cusp 4-5 | Cusp 4-6 | Cusp 5-6 |
| --- | --- | --- | --- | --- | --- | --- | --- | --- | --- | --- | --- | --- | --- | --- | --- | --- | --- |
| dm_2_ | Cusp 5 | Coefficient | - | - | - | - | - | - | - | - | - | - | - | - | - | - | - |
|  |  | p-value^1^ | - | - | - | - | - | - | - | - | - | - | - | - | - | - | - |
|  |  | Odds ratio^2^ | - | - | - | - | - | - | - | - | - | - | - | - | - | - | - |
|  | Cusp 6 | Coefficient | -0.24 | 0.76 | 0.23 | 0.34 | - | 0.36 | 0.06 | 0.58 | - | 0.82 | -0.67 | - | **2.50** | - | - |
|  |  | p-value | 0.64 | 0.14 | 0.54 | 0.40 | - | 0.35 | 0.87 | 0.17 | - | 0.12 | 0.33 | - | **0.00** | - | - |
|  |  | Odds ratio | - | - | - | - | - | - | - | - | - | - | - | - | **1.28** | - | - |
|  | Cusp 7 | Coefficient | -0.55 | 0.36 | -0.18 | 0.01 | -0.35 | -0.35 | -0.36 | -0.38 | -0.69 | -0.75 | -1.09 | -0.84 | -0.06 | -1.22 | -0.35 |
|  |  | p-value | 0.31 | 0.46 | 0.63 | 0.98 | 0.45 | 0.37 | 0.38 | 0.35 | 0.21 | 0.16 | 0.14 | 0.23 | 0.91 | 0.14 | 0.63 |
|  |  | Odds ratio | - | - | - | - | - | - | - | - | - | - | - | - | - | - | - |
| LM_1_ | Cusp 5 | Coefficient | -0.18 | **-2.01** | 0.02 | - | - | -0.45 | -0.15 | - | - | **1.36** | - | - | - | - | - |
|  |  | p-value | 0.78 | **0.01** | 0.97 | - | - | 0.40 | 0.76 | - | - | **0.04** | - | - | - | - | - |
|  |  | Odds ratio | - | **0.82** | - | - | - | - | - | - | - | **1.15** | - | - | - | - | - |
|  | Cusp 6 | Coefficient | -0.05 | -0.14 | -0.15 | -0.15 | **-** | 0.18 | -0.27 | 0.27 | **-** | **0.54** | -0.36 | **-** | **1.21** | - | - |
|  |  | p-value | 0.86 | 0.58 | 0.46 | 0.53 | **-** | 0.38 | 0.20 | 0.22 | **-** | **0.03** | 0.24 | **-** | **0.00** | - | - |
|  |  | Odds ratio | - | - | - | - | - | - | - | - | - | **1.06** | - | - | **1.13** | - | - |
|  | Cusp 7 | Coefficient | -0.01 | -0.24 | 0.11 | 0.01 | -0.50 | -0.25 | 0.37 | -0.04 | -0.42 | -0.25 | 0.05 | -1.04 | -0.43 | -0.54 | -0.09 |
|  |  | p-value | 0.98 | 0.44 | 0.66 | 0.96 | 0.24 | 0.34 | 0.17 | 0.88 | 0.25 | 0.38 | 0.89 | 0.10 | 0.13 | 0.31 | 0.87 |
|  |  | Odds ratio | - | - | - | - | - | - | - | - | - | - | - | - | - | - | - |
| RM_1_ | Cusp 5 | Coefficient | -0.03 | **-3.15** | -0.09 | - | - | -0.33 | 0.08 | - | - | **1.89** | - | - | - | - | - |
|  |  | p-value | 0.97 | **0.00** | 0.84 | - | - | 0.53 | 0.86 | - | - | **0.00** | - | - | - | - | - |
|  |  | Odds ratio | - | **0.73** | - | - | - | - | - | - | - | **1.21** | - | - | - | - | - |
|  | Cusp 6 | Coefficient | -0.16 | -0.18 | -0.11 | -0.14 | - | 0.37 | -0.33 | 0.39 | - | **0.89** | -0.46 | - | **1.61** | - | - |
|  |  | p-value | 0.62 | 0.54 | 0.62 | 0.57 | - | 0.16 | 0.17 | 0.12 | - | **0.00** | 0.16 | - | **0.00** | - | - |
|  |  | Odds ratio | - | - | - | - | - | - | - | - | - | **1.09** | - | - | **1.18** | - | - |
|  | Cusp 7 | Coefficient | 0.08 | -0.13 | 0.32 | 0.27 | 0.32 | -0.16 | **0.70** | 0.01 | 0.32 | -0.22 | 0.59 | 0.02 | **-0.79** | **-0.44** | -0.17 |
|  |  | p-value | 0.85 | 0.72 | 0.28 | 0.39 | 0.64 | 0.64 | **0.03** | 0.98 | 0.62 | 0.50 | 0.15 | 0.97 | **0.02** | **0.44** | 0.80 |
|  |  | Odds ratio | - | - | - | - | - | - | **1.07** | - | - | - | - | - | **0.92** | **0.96** | - |

**S12 Table. Results of logistic regression when using AICD as a predictor of lower molar accessory cusp development (morphology dichotomized).** Cells highlighted in blue represent intercusp distances that become significantly longer with accessory cusp development. Cells highlighted in red represent intercusp distances that become significantly shorter with accessory cusp development.

^1^α≤0.05; significant results in bold

^2^Odds ratio obtained by dividing the model coefficient by 10 to scale the odds ratio to a 0.10-unit change and exponentiating the quotient

|  | | | Cusp 1-2 | Cusp 1-3 | Cusp 1-4 | Cusp 1-5 | Cusp 1-6 | Cusp 2-3 | Cusp 2-4 | Cusp 2-5 | Cusp 2-6 | Cusp 3-4 | Cusp 3-5 | Cusp 3-6 | Cusp 4-5 | Cusp 4-6 | Cusp 5-6 |
| --- | --- | --- | --- | --- | --- | --- | --- | --- | --- | --- | --- | --- | --- | --- | --- | --- | --- |
| dm_2_ | Cusp 5 | Coefficient | - | - | - | - | - | - | - | - | - | - | - | - | - | - | - |
|  |  | p-value^1^ | - | - | - | - | - | - | - | - | - | - | - | - | - | - | - |
|  |  | Odds ratio^2^ | - | - | - | - | - | - | - | - | - | - | - | - | - | - | - |
|  | Cusp 6 | Coefficient | -5.34 | 5.75 | -0.08 | 1.12 | - | 2.17 | -1.52 | 5.15 | - | 6.58 | -13.04 | - | **30.45** | - | - |
|  |  | p-value | 0.33 | 0.25 | 0.99 | 0.80 | - | 0.61 | 0.70 | 0.33 | - | 0.26 | 0.11 | - | **0.00** | - | - |
|  |  | Odds ratio | - | - | - | - | - | - | - | - | - | - | - | - | **21.00** | - | - |
|  | Cusp 7 | Coefficient | -5.97 | 4.40 | -1.77 | 0.49 | -5.73 | -4.47 | -3.69 | -6.55 | -13.26 | -10.36 | -13.64 | -13.45 | -0.31 | -13.37 | -3.46 |
|  |  | p-value | 0.30 | 0.38 | 0.66 | 0.92 | 0.34 | 0.32 | 0.37 | 0.23 | 0.11 | 0.10 | 0.10 | 0.15 | 0.96 | 0.12 | 0.63 |
|  |  | Odds ratio | - | - | - | - | - | - | - | - | - | - | - | - | - | - | - |
| LM_1_ | Cusp 5 | Coefficient | **-22.68** | **-29.97** | **-19.80** | - | - | **-27.24** | -14.80 | - | - | 8.42 | - | - | - | - | - |
|  |  | p-value | **0.03** | **0.00** | **0.02** | - | - | **0.00** | 0.07 | - | - | 0.30 | - | - | - | - | - |
|  |  | Odds ratio | **0.10** | **0.05** | **0.14** | - | - | **0.07** | - | - | - | - | - | - | - | - | - |
|  | Cusp 6 | Coefficient | -4.01 | -3.43 | **-6.06** | -4.43 | **-** | 0.15 | **-5.44** | 3.66 | **-** | **8.10** | -5.53 | **-** | **19.53** | - | - |
|  |  | p-value | 0.25 | 0.21 | **0.04** | 0.16 | **-** | 0.96 | **0.04** | 0.28 | **-** | **0.03** | 0.12 | **-** | **0.00** | - | - |
|  |  | Odds ratio | - | - | **0.55** | - | - | - | **0.58** | - | - | **2.25** | - | - | **7.05** | - | - |
|  | Cusp 7 | Coefficient | -3.90 | -5.23 | -1.74 | -2.97 | -13.44 | **-7.77** | 2.58 | -5.29 | -7.93 | **-9.99** | -1.14 | -15.55 | **-8.52** | -6.94 | 1.78 |
|  |  | p-value | 0.37 | 0.15 | 0.62 | 0.42 | 0.08 | **0.03** | 0.41 | 0.19 | 0.16 | **0.03** | 0.79 | 0.05 | **0.03** | 0.29 | 0.80 |
|  |  | Odds ratio | - | - | - | - | - | **0.46** | - | - | - | **0.37** | - | - | **0.43** | - | - |
| RM_1_ | Cusp 5 | Coefficient | **-30.77** | **-56.28** | **-30.36** | - | - | **-24.82** | -9.56 | - | - | 15.21 | - | - | - | - | - |
|  |  | p-value | **0.02** | **0.01** | **0.01** | - | - | **0.00** | 0.13 | - | - | 0.07 | - | - | - | - | - |
|  |  | Odds ratio | **0.05** | **0.00** | **0.05** | - | - | **0.08** | - | - | - | - | - | - | - | - | - |
|  | Cusp 6 | Coefficient | -5.88 | -3.58 | -4.58 | -3.13 | - | 2.04 | **-5.45** | 6.13 | - | **16.36** | -6.79 | - | **25.32** | - | - |
|  |  | p-value | 0.15 | 0.24 | 0.12 | 0.33 | - | 0.53 | **0.05** | 0.09 | - | **0.00** | 0.09 | - | **0.00** | - | - |
|  |  | Odds ratio | - | - | - | - | - | - | **0.58** | - | - | **5.13** | - | - | **12.58** | - | - |
|  | Cusp 7 | Coefficient | -4.39 | -4.48 | 0.13 | -0.01 | -1.03 | -7.84 | 5.22 | -5.47 | -0.46 | **-11.85** | 5.17 | -2.87 | **-15.42** | -7.51 | -4.58 |
|  |  | p-value | 0.37 | 0.28 | 0.97 | 1 | 0.90 | 0.07 | 0.14 | 0.20 | 0.95 | **0.02** | 0.30 | 0.23 | **0.00** | 0.28 | 0.25 |
|  |  | Odds ratio | - | - | - | - | - | - | - | - | - | **0.31** | - | - | **0.21** | - | - |

**S13 Table. Results of logistic regression when using RICD scaled by BLxMD crown area as a predictor of lower molar accessory cusp development (morphology dichotomized).** Cells highlighted in blue represent intercusp distances that become significantly longer with accessory cusp development. Cells highlighted in red represent intercusp distances that become significantly shorter with accessory cusp development.

^1^α≤0.05; significant results in bold

^2^Odds ratio obtained by dividing the model coefficient by 10 to scale the odds ratio to a 0.10-unit change and exponentiating the quotient

|  | | | Cusp 1-2 | Cusp 1-3 | Cusp 1-4 | Cusp 1-5 | Cusp 1-6 | Cusp 2-3 | Cusp 2-4 | Cusp 2-5 | Cusp 2-6 | Cusp 3-4 | Cusp 3-5 | Cusp 3-6 | Cusp 4-5 | Cusp 4-6 | Cusp 5-6 |
| --- | --- | --- | --- | --- | --- | --- | --- | --- | --- | --- | --- | --- | --- | --- | --- | --- | --- |
| dm_2_ | Cusp 5 | Coefficient | - | - | - | - | - | - | - | - | - | - | - | - | - | - | - |
|  |  | p-value^1^ | - | - | - | - | - | - | - | - | - | - | - | - | - | - | - |
|  |  | Odds ratio^2^ | - | - | - | - | - | - | - | - | - | - | - | - | - | - | - |
|  | Cusp 6 | Coefficient | -4.26 | 5.87 | 0.53 | 1.85 | - | 2.63 | -0.92 | 5.43 | - | 6.70 | -9.92 | - | **26.70** | - | - |
|  |  | p-value | 0.39 | 0.21 | 0.88 | 0.65 | - | 0.50 | 0.80 | 0.24 | - | 0.20 | 0.16 | - | **0.00** | - | - |
|  |  | Odds ratio | - | - | - | - | - | - | - | - | - | - | - | - | **14.44** | - | - |
|  | Cusp 7 | Coefficient | -5.39 | 4.16 | -1.47 | 0.46 | -5.75 | -4.06 | -3.22 | -5.45 | -12.82 | -8.75 | -11.96 | -14.07 | -0.42 | -12.31 | -3.67 |
|  |  | p-value | 0.30 | 0.36 | 0.68 | 0.91 | 0.30 | 0.32 | 0.38 | 0.24 | 0.09 | 0.11 | 0.11 | 0.13 | 0.94 | 0.11 | 0.57 |
|  |  | Odds ratio | - | - | - | - | - | - | - | - | - | - | - | - | - | - | - |
| LM_1_ | Cusp 5 | Coefficient | -13.48 | **-28.03** | -11.15 | - | - | **-18.54** | -8.68 | - | - | 12.40 | - | - | - | - | - |
|  |  | p-value | 0.12 | **0.00** | 0.15 | - | - | **0.02** | 0.19 | - | - | 0.10 | - | - | - | - | - |
|  |  | Odds ratio | - | **0.06** | - | - | - | **0.16** | - | - | - | - | - | - | - | - | - |
|  | Cusp 6 | Coefficient | -2.47 | -2.51 | -4.22 | -3.11 | **-** | 1.19 | -4.24 | 3.67 | **-** | **7.98** | -4.55 | **-** | **18.27** | - | - |
|  |  | p-value | 0.44 | 0.33 | 0.11 | 0.27 | **-** | 0.63 | 0.07 | 0.21 | **-** | **0.01** | 0.15 | **-** | **0.00** | - | - |
|  |  | Odds ratio | - | - | - | - | - | - | - | - | - | **2.22** | - | - | **6.21** | - | - |
|  | Cusp 7 | Coefficient | -4.44 | -5.35 | -2.47 | -3.80 | -12.73 | **-7.53** | 1.85 | -5.74 | -7.19 | **-9.27** | -1.88 | **-16.00** | **-8.15** | -6.76 | -0.33 |
|  |  | p-value | 0.27 | 0.11 | 0.44 | 0.25 | 0.06 | **0.02** | 0.52 | 0.10 | 0.14 | **0.02** | 0.62 | **0.04** | **0.02** | 0.26 | 0.96 |
|  |  | Odds ratio | - | - | - | - | - | **0.47** | - | - | - | **0.40** | - | **0.20** | **0.44** | - | - |
| RM_1_ | Cusp 5 | Coefficient | -17.75 | **-47.17** | **-17.35** | - | - | **-19.57** | -5.76 | - | - | **19.34** | - | - | - | - | - |
|  |  | p-value | 0.08 | **0.00** | **0.04** | - | - | **0.01** | 0.30 | - | - | **0.02** | - | - | - | - | - |
|  |  | Odds ratio | - | **0.01** | **0.18** | - | - | **0.14** | - | - | - | **6.92** | - | - | - | - | - |
|  | Cusp 6 | Coefficient | -4.52 | -3.00 | -3.42 | -2.52 | - | 3.05 | -4.67 | 5.88 | - | **15.41** | -5.88 | - | **22.27** | - | - |
|  |  | p-value | 0.23 | 0.29 | 0.21 | 0.39 | - | 0.33 | 0.06 | 0.07 | - | **0.00** | 0.11 | - | **0.00** | - | - |
|  |  | Odds ratio | - | - | - | - | - | - | - | - | - | **4.67** | - | - | **9.28** | - | - |
|  | Cusp 7 | Coefficient | -4.51 | -4.58 | -0.20 | -0.42 | -4.49 | **-8.33** | 4.76 | -5.35 | -3.28 | **-10.77** | 4.29 | -2.83 | **-14.00** | -5.01 | -4.53 |
|  |  | p-value | 0.32 | 0.25 | 0.95 | 0.91 | 0.55 | **0.05** | 0.15 | 0.17 | 0.64 | **0.02** | 0.34 | 0.20 | **0.00** | 0.12 | 0.21 |
|  |  | Odds ratio | - | - | - | - | - | **0.43** | - | - | - | **0.34** | - | - | **0.25** | - | - |

**S14 Table. Results of logistic regression when using RICD scaled by trace crown area as a predictor of lower molar accessory cusp development (morphology dichotomized).** Cells highlighted in blue represent intercusp distances that become significantly longer with accessory cusp development. Cells highlighted in red represent intercusp distances that become significantly shorter with accessory cusp development.

^1^α≤0.05; significant results in bold

^2^Odds ratio obtained by dividing the model coefficient by 10 to scale the odds ratio to a 0.10-unit change and exponentiating the quotient

|  | | | Cusp 1-2 | Cusp 1-3 | Cusp 1-4 | Cusp 1-5 | Cusp 1-6 | Cusp 2-3 | Cusp 2-4 | Cusp 2-5 | Cusp 2-6 | Cusp 3-4 | Cusp 3-5 | Cusp 3-6 | Cusp 4-5 | Cusp 4-6 | Cusp 5-6 |
| --- | --- | --- | --- | --- | --- | --- | --- | --- | --- | --- | --- | --- | --- | --- | --- | --- | --- |
| dm_2_ | Cusp 5 | Coefficient | - | - | - | - | - | - | - | - | - | - | - | - | - | - | - |
|  |  | p-value^1^ | - | - | - | - | - | - | - | - | - | - | - | - | - | - | - |
|  |  | Odds ratio^2^ | - | - | - | - | - | - | - | - | - | - | - | - | - | - | - |
|  | Cusp 6 | Coefficient | -1.55 | 2.56 | 0.36 | 0.90 | - | 1.01 | -0.27 | 2.20 | - | 2.76 | -3.65 | - | **10.53** | - | - |
|  |  | p-value | 0.40 | 0.17 | 0.81 | 0.59 | - | 0.48 | 0.85 | 0.22 | - | 0.18 | 0.16 | - | **0.00** | - | - |
|  |  | Odds ratio | - | - | - | - | - | - | - | - | - | - | - | - | **2.87** | - | - |
|  | Cusp 7 | Coefficient | -2.07 | 1.65 | -0.76 | 0.18 | -1.50 | -1.44 | -1.37 | -1.97 | -3.57 | -3.40 | -4.38 | -2.97 | -0.02 | -4.39 | -0.74 |
|  |  | p-value | 0.30 | 0.36 | 0.62 | 0.92 | 0.46 | 0.34 | 0.36 | 0.27 | 0.17 | 0.12 | 0.11 | 0.24 | 0.99 | 0.14 | 0.75 |
|  |  | Odds ratio | - | - | - | - | - | - | - | - | - | - | - | - | - | - | - |
| LM_1_ | Cusp 5 | Coefficient | -3.31 | **-9.66** | -2.35 | - | - | **-4.79** | -2.36 | - | - | 4.38 | - | - | - | - | - |
|  |  | p-value | 0.22 | **0.00** | 0.28 | - | - | **0.05** | 0.27 | - | - | 0.08 | - | - | - | - | - |
|  |  | Odds ratio | - | **0.38** | - | - | - | **0.62** | - | - | - | - | - | - | - | - | - |
|  | Cusp 6 | Coefficient | -0.50 | -0.72 | -1.04 | -0.82 | - | 0.62 | -1.30 | 1.50 | - | **2.52** | -1.44 | - | **5.40** | - | - |
|  |  | p-value | 0.64 | 0.43 | 0.23 | 0.40 | - | 0.46 | 0.11 | 0.12 | - | **0.01** | 0.19 | - | **0.00** | - | - |
|  |  | Odds ratio | - | - | - | - | - | - | - | - | - | **1.29** | - | - | **1.72** | - | - |
|  | Cusp 7 | Coefficient | -0.61 | -1.33 | 0.03 | -0.43 | -3.19 | -1.69 | 1.20 | -0.78 | -2.21 | -1.83 | -0.10 | -4.96 | -2.00 | -1.99 | -0.21 |
|  |  | p-value | 0.65 | 0.25 | 0.98 | 0.71 | 0.13 | 0.11 | 0.25 | 0.50 | 0.17 | 0.13 | 0.94 | 0.06 | 0.07 | 0.31 | 0.92 |
|  |  | Odds ratio | - | - | - | - | - | - | - | - | - | - | - | - | - | - | - |
| RM_1_ | Cusp 5 | Coefficient | -3.88 | **-15.11** | -3.97 | - | - | **-4.98** | -1.47 | - | - | **6.48** | - | - | - | - | - |
|  |  | p-value | 0.18 | **0.00** | 0.09 | - | - | **0.03** | 0.43 | - | - | **0.02** | - | - | - | - | - |
|  |  | Odds ratio | - | **0.22** | - | - | - | **0.61** | - | - | - | **1.91** | - | - | - | - | - |
|  | Cusp 6 | Coefficient | -1.07 | -0.84 | -0.84 | -0.65 | - | 1.36 | -1.49 | **2.40** | - | **4.42** | -1.89 | - | **7.07** | - | - |
|  |  | p-value | 0.39 | 0.40 | 0.37 | 0.53 | - | 0.19 | 0.09 | **0.04** | - | **0.00** | 0.13 | - | **0.00** | - | - |
|  |  | Odds ratio | - | - | - | - | - | - | - | **1.27** | - | **1.56** | - | - | **2.03** | - | - |
|  | Cusp 7 | Coefficient | -0.24 | -0.83 | 1.09 | 0.99 | 1.84 | -1.31 | **2.58** | -0.48 | 1.60 | -1.69 | 2.21 | 0.05 | **-3.56** | -1.50 | -0.64 |
|  |  | p-value | 0.88 | 0.53 | 0.37 | 0.46 | 0.57 | 0.32 | **0.04** | 0.72 | 0.57 | 0.22 | 0.16 | 0.98 | **0.01** | 0.46 | 0.78 |
|  |  | Odds ratio | - | - | - | - | - | - | **1.29** | - | - | - | - | - | **0.70** | - | - |

**S15 Table. Results of logistic regression when using RICD scaled by mesiodistal tooth dimension as a predictor of lower molar accessory cusp development (morphology dichotomized).** Cells highlighted in blue represent intercusp distances that become significantly longer with accessory cusp development. Cells highlighted in red represent intercusp distances that become significantly shorter with accessory cusp development.

^1^α≤0.05; significant results in bold

^2^Odds ratio obtained by dividing the model coefficient by 10 to scale the odds ratio to a 0.10-unit change and exponentiating the quotient

**S16 Table. Results of logistic regression when using RICD scaled by buccolingual tooth dimension as a predictor of lower molar accessory cusp development (morphology dichotomized).** Cells highlighted in blue represent intercusp distances that become significantly longer with accessory cusp development. Cells highlighted in red represent intercusp distances that become significantly shorter with accessory cusp development.

|  | | | Cusp 1-2 | Cusp 1-3 | Cusp 1-4 | Cusp 1-5 | Cusp 1-6 | Cusp 2-3 | Cusp 2-4 | Cusp 2-5 | Cusp 2-6 | Cusp 3-4 | Cusp 3-5 | Cusp 3-6 | Cusp 4-5 | Cusp 4-6 | Cusp 5-6 |
| --- | --- | --- | --- | --- | --- | --- | --- | --- | --- | --- | --- | --- | --- | --- | --- | --- | --- |
| dm_2_ | Cusp 5 | Coefficient | - | - | - | - | - | - | - | - | - | - | - | - | - | - | - |
|  |  | p-value^1^ | - | - | - | - | - | - | - | - | - | - | - | - | - | - | - |
|  |  | Odds ratio^2^ | - | - | - | - | - | - | - | - | - | - | - | - | - | - | - |
|  | Cusp 6 | Coefficient | -0.91 | 1.87 | 0.40 | 0.72 | - | 0.92 | -0.03 | 1.62 | - | 2.27 | -2.55 | - | **8.22** | - | - |
|  |  | p-value | 0.54 | 0.19 | 0.70 | 0.53 | - | 0.43 | 0.98 | 0.21 | - | 0.16 | 0.24 | - | **0.00** | - | - |
|  |  | Odds ratio | - | - | - | - | - | - | - | - | - | - | - | - | **2.27** | - | - |
|  | Cusp 7 | Coefficient | -1.65 | 1.00 | -0.50 | 0.02 | -1.39 | -1.14 | -1.03 | -1.34 | -2.67 | -2.51 | -3.59 | -3.92 | -0.28 | -3.84 | -1.54 |
|  |  | p-value | 0.31 | 0.46 | 0.64 | 0.98 | 0.33 | 0.34 | 0.38 | 0.30 | 0.13 | 0.13 | 0.12 | 0.13 | 0.86 | 0.12 | 0.50 |
|  |  | Odds ratio | - | - | - | - | - | - | - | - | - | - | - | - | - | - | - |
| LM_1_ | Cusp 5 | Coefficient | -2.85 | **-7.54** | -1.78 | - | - | -4.05 | -1.82 | - | - | 3.87 | - | - | - | - | - |
|  |  | p-value | 0.25 | **0.00** | 0.34 | - | - | 0.06 | 0.32 | - | - | 0.08 | - | - | - | - | - |
|  |  | Odds ratio | - | **0.47** | - | - | - | - | - | - | - | - | - | - | - | - | - |
|  | Cusp 6 | Coefficient | -0.77 | -0.83 | -1.14 | -0.97 | - | 0.25 | -1.27 | 0.74 | - | **1.90** | -1.46 | - | **4.71** | - | - |
|  |  | p-value | 0.42 | 0.30 | 0.13 | 0.24 | - | 0.74 | 0.07 | 0.35 | - | **0.04** | 0.14 | - | **0.00** | - | - |
|  |  | Odds ratio | - | - | - | - | - | - | - | - | - | **1.21** | - | - | **1.60** | - | - |
|  | Cusp 7 | Coefficient | -0.52 | -1.12 | 0.03 | -0.32 | -2.13 | -1.39 | 0.98 | -0.60 | -1.57 | -1.62 | -0.06 | -3.98 | -1.95 | -1.79 | -0.12 |
|  |  | p-value | 0.67 | 0.27 | 0.97 | 0.74 | 0.19 | 0.13 | 0.27 | 0.53 | 0.24 | 0.14 | 0.96 | 0.08 | 0.06 | 0.32 | 0.95 |
|  |  | Odds ratio | - | - | - | - | - | - | - | - | - | - | - | - | - | - | - |
| RM_1_ | Cusp 5 | Coefficient | -2.40 | **-11.91** | -2.27 | - | - | -3.66 | -0.82 | - | - | **5.73** | - | - | - | - | - |
|  |  | p-value | 0.32 | **0.00** | 0.21 | - | - | 0.07 | 0.60 | - | - | **0.01** | - | - | - | - | - |
|  |  | Odds ratio | - | **0.30** | - | - | - | - | - | - | - | **1.77** | - | - | - | - | - |
|  | Cusp 6 | Coefficient | -1.19 | -0.92 | -0.87 | -0.74 | - | 0.85 | -1.35 | 1.21 | - | **3.52** | -1.74 | - | **6.23** | - | - |
|  |  | p-value | 0.28 | 0.31 | 0.26 | 0.37 | - | 0.35 | 0.08 | 0.17 | - | **0.00** | 0.11 | - | **0.00** | - | - |
|  |  | Odds ratio | - | - | - | - | - | - | - | - | - | **1.42** | - | - | **1.86** | - | - |
|  | Cusp 7 | Coefficient | -0.66 | -1.02 | 0.44 | 0.29 | -0.12 | -1.52 | 1.70 | -0.78 | 0.02 | -1.87 | 1.53 | -0.59 | **-3.62** | -2.02 | -0.87 |
|  |  | p-value | 0.63 | 0.40 | 0.66 | 0.78 | 0.96 | 0.19 | 0.09 | 0.46 | 0.99 | 0.13 | 0.26 | 0.78 | **0.00** | 0.31 | 0.67 |
|  |  | Odds ratio | - | - | - | - | - | - | - | - | - | - | - | - | **0.70** | - | - |

^1^α≤0.05; significant results in bold

^2^Odds ratio obtained by dividing the model coefficient by 10 to scale the odds ratio to a 0.10-unit change and exponentiating the quotient

|  | | | AICA1-4^4^ | AICA1-5^5^ | AICA1-6^6^ | RICA1-4 | RICA1-5 | RICA1-6 | TICA1-4 | TICA1-5 | TICA1-6 | MDICA1-4 | MDICA1-5 | MDICA1-6 | BLICA1-4 | BLICA1-5 | BLICA1-6 |
| --- | --- | --- | --- | --- | --- | --- | --- | --- | --- | --- | --- | --- | --- | --- | --- | --- | --- |
| dm_2_ | Cusp 5 | Coefficient | 0.12 | - | - | 1.41 | - | - | 1.35 | - | - | 0.47 | - | - | 0.36 | - | - |
|  |  | LRχ^1^ | 2.52 | - | - | 3.04 | - | - | 3.38 | - | - | 2.90 | - | - | 2.65 | - | - |
|  |  | p-value^2^ | 0.11 | - | - | 0.08 | - | - | 0.07 | - | - | 0.09 | - | - | 0.10 | - | - |
|  |  | Odds ratio^3^ | - | - | - | - | - | - | - | - | - | - | - | - | - | - | - |
|  | Cusp 6 | Coefficient | **-0.21** | **-0.13** | **-** | **-2.37** | **-1.51** | - | **-2.16** | **-1.39** | - | **-0.79** | **-0.49** | - | **-0.63** | **-0.39** | - |
|  |  | LRχ | **6.60** | **4.15** | **-** | **7.54** | **4.78** | - | **7.73** | **5.00** | - | **7.23** | **4.48** | - | **6.99** | **4.49** | - |
|  |  | p-value | **0.01** | **0.04** | **-** | **0.01** | **0.03** | - | **0.01** | **0.03** | - | **0.01** | **0.03** | - | **0.01** | **0.03** | - |
|  |  | Odds ratio | **0.98** | **0.99** | - | **0.79** | **0.86** | - | **0.81** | **0.87** | - | **0.92** | **0.95** | - | **0.94** | **0.96** | - |
|  | Cusp 7 | Coefficient | 0.02 | -0.05* | -0.06* | -0.05* | -0.86* | -0.92* | -0.16* | -0.87* | -0.93* | 0.08 | -0.18 | -0.20* | -0.03* | -0.24* | -0.25* |
|  |  | LRχ | 0.01 | 0.22 | 0.31 | 0.00 | 0.55 | 0.73 | 0.02 | 0.71 | 0.93 | 0.03 | 0.20 | 0.29 | 0.01 | 0.53 | 0.71 |
|  |  | p-value | 0.90 | 0.64 | 0.58 | 0.97 | 0.46 | 0.39 | 0.90 | 0.40 | 0.33 | 0.85 | 0.65 | 0.59 | 0.94 | 0.47 | 0.40 |
|  |  | Odds ratio | - | - | - | - | - | - | - | - | - | - | - | - | - | - | - |
| LM_1_ | Cusp 5 | Coefficient | -0.00 | **-** | - | -0.21 | - | - | -0.18 | - | - | -0.05 | - | - | -0.01 | - | - |
|  |  | LRχ | 0.01 | **-** | - | 0.24 | - | - | 0.23 | - | - | 0.19 | - | - | 0.01 | - | - |
|  |  | p-value | 0.94 | **-** | - | 0.62 | - | - | 0.63 | - | - | 0.66 | - | - | 0.93 | - | - |
|  |  | Odds ratio | - | **-** | - | - | - | - | - | - | - | - | - | - | - | - | - |
|  | Cusp 6 | Coefficient | -0.00 | -0.00 | - | -0.25 | -0.25 | - | -0.17 | -0.17 | **-** | -0.02 | -0.01 | - | -0.07 | -0.06 | - |
|  |  | LRχ | 0.01 | 0.01 | - | 0.32 | 0.39 | - | 0.19 | 0.22 | **-** | 0.02 | 0.01 | - | 0.30 | 0.35 | - |
|  |  | p-value | 0.91 | 0.94 | - | 0.57 | 0.53 | - | 0.66 | 0.64 | **-** | 0.89 | 0.93 | - | 0.58 | 0.56 | - |
|  |  | Odds ratio | - | - | - | - | - | - | - | - | - | - | - | - | - | - | - |
|  | Cusp 7 | Coefficient | -0.08 | -0.08 | -0.07 | -1.14 | **-1.17** | **-1.31** | -1.05 | **-1.07** | **-1.19** | -0.32 | **-0.32** | **-0.32** | -0.30 | **-0.30** | -0.33 |
|  |  | LRχ | 2.82 | 3.59 | 3.70 | 3.41 | **4.84** | **5.56** | 3.70 | **5.18** | **5.98** | 3.06 | **4.11** | **4.36** | 3.34 | **4.47** | 1.54 |
|  |  | p-value | 0.09 | 0.06 | 0.05 | 0.06 | **0.03** | **0.02** | 0.05 | **0.02** | **0.01** | 0.08 | **0.04** | **0.04** | 0.07 | **0.03** | 0.21 |
|  |  | Odds ratio | - | - | - | - | **0.89** | **0.88** | - | **0.90** | **0.89** | - | **0.97** | **0.97** | - | **0.97** | - |
| RM_1_ | Cusp 5 | Coefficient | 0.03 | **-** | - | 0.23 | - | **-** | 0.28 | - | **-** | 0.06 | - | - | 0.12 | - | - |
|  |  | LRχ | 0.56 | **-** | - | 0.21 | - | **-** | 0.34 | - | **-** | 0.17 | - | - | 0.68 | - | - |
|  |  | p-value | 0.46 | **-** | - | 0.65 | - | **-** | 0.56 | - | **-** | 0.68 | - | - | 0.41 | - | - |
|  |  | Odds ratio | - | - | - | - | - | - | - | - | - | - | - | - | - | - | - |
|  | Cusp 6 | Coefficient | -0.05 | -0.04 | - | **-1.06** | **-1.15** | - | **-0.96** | **-1.02** | - | -0.22 | -0.21 | - | -0.26 | **-0.26** | - |
|  |  | LRχ | 1.33 | 1.29 | - | **4.23** | **5.61** | - | **4.06** | **5.22** | - | 1.91 | 2.15 | - | 3.34 | **3.92** | - |
|  |  | p-value | 0.25 | 0.26 | - | **0.04** | **0.02** | - | **0.04** | **0.02** | - | 0.17 | 0.14 | - | 0.07 | **0.05** | - |
|  |  | Odds ratio | - | - | - | **0.90** | **0.89** | - | **0.91** | **0.90** | - | - | - |  | - | **0.97** |  |
|  | Cusp 7 | Coefficient | -0.00 | -0.02 | -0.03 | -0.18 | -0.52 | -0.70 | -0.29 | -0.61 | -0.81 | -0.00 | -0.10 | -0.13 | -0.04 | -0.13 | -0.16 |
|  |  | LRχ | 0.00 | 0.18 | 0.26 | 0.05 | 0.55 | 0.87 | 0.16 | 0.89 | 1.38 | 0.00 | 0.23 | 0.36 | 0.04 | 0.45 | 0.66 |
|  |  | p-value | 0.99 | 0.67 | 0.61 | 0.82 | 0.46 | 0.35 | 0.69 | 0.34 | 0.24 | 0.99 | 0.63 | 0.55 | 0.84 | 0.50 | 0.42 |
|  |  | Odds ratio | - | - | - | - | - | - | - | - | - | - | - | - | - | - | - |

**S17 Table. Results of proportional ordinal regression when using intercusp area as a predictor of lower molar accessory cusp trait expression.** Cells highlighted in blue represent intercusp areas that become significantly larger with increased accessory cusp trait expression. Cells highlighted in red represent intercusp areas that become significantly smaller with increased accessory cusp trait expression.

^*^Values violating the assumption of proportionality are marked with an asterisk, these results should be interpreted with caution

^1^LRχ: Likelihood ratio chi-square test statistic

^2^α≤0.05; significant results in bold

^3^Odds ratio obtained by dividing the model coefficient by 10 to scale the odds ratio to a 0.10-unit change and exponentiating the quotient

^4^Absolute intercusp area formed by protoconid-metaconid-hypoconid-entoconid polygon

^5^Absolute intercusp area formed by protoconid-metaconid-hypoconid-entoconid-hypoconulid polygon

^6^Absolute intercusp area formed by protoconid-metaconid-hypoconid-entoconid-hypoconulid-cusp6 polygon

|  | | | AICA1-4^3^ | AICA1-5^4^ | AICA1-6^5^ | RICA1-4 | RICA1-5 | RICA1-6 | TICA1-4 | TICA1-5 | TICA1-6 | MDICA1-4 | MDICA1-5 | MDICA1-6 | BLICA1-4 | BLICA1-5 | BLICA1-6 |
| --- | --- | --- | --- | --- | --- | --- | --- | --- | --- | --- | --- | --- | --- | --- | --- | --- | --- |
| dm_2_ | Cusp 5 | Coefficient | - | - | - | - | - | - | - | - | - | - | - | - | - | - | - |
|  |  | p-value^1^ | - | - | - | - | - | - | - | - | - | - | - | - | - | - | - |
|  |  | Odds ratio^2^ | - | - | - | - | - | - | - | - | - | - | - | - | - | - | - |
|  | Cusp 6 | Coefficient | -0.34 | -0.31 | **-** | -3.35 | -3.48 | - | -2.94 | -3.48 | - | -1.14 | -1.10 | - | -0.99 | -0.92 | - |
|  |  | p-value | 0.21 | 0.19 | **-** | 0.21 | 0.19 | - | 0.21 | 0.21 | - | 0.21 | 0.19 | - | 0.20 | 0.18 | - |
|  |  | Odds ratio | - | - | - | - | - | - | - | - | - | - | - | - | - | - | - |
|  | Cusp 7 | Coefficient | -0.01 | -0.07 | -0.08 | -0.37 | -1.23 | -1.37 | -0.46 | -1.22 | -1.37 | -0.01 | -0.27 | -0.30 | -0.12 | -0.33 | -0.36 |
|  |  | p-value | 0.94 | 0.53 | 0.46 | 0.79 | 0.35 | 0.29 | 0.72 | 0.30 | 0.24 | 0.98 | 0.53 | 0.46 | 0.77 | 0.37 | 0.30 |
|  |  | Odds ratio | - | - | - | - | - | - | - | - | - | - | - | - | - | - | - |
| LM_1_ | Cusp 5 | Coefficient | -0.04 | - | - | -1.58 | - | - | -1.09 | - | - | -0.29 | - | - | -0.02 | - | - |
|  |  | p-value | 0.51 | - | - | 0.07 | - | - | 0.15 | - | - | 0.22 | - | - | 0.77 | - | - |
|  |  | Odds ratio | - | - | - | - | - | - | - | - | - | - | - | - | - | - | - |
|  | Cusp 6 | Coefficient | 0.00 | 0.01 | **-** | -0.14 | 0.15 | - | -0.06 | 0.17 | - | -0.01 | 0.06 | - | -0.02 | 0.03 | - |
|  |  | p-value | 0.97 | 0.51 | **-** | 0.64 | 0.59 | - | 0.82 | 0.51 | - | 0.93 | 0.46 | - | 0.77 | 0.63 | - |
|  |  | Odds ratio | - | - | - | - | - | - | - | - | - | - | - | - | - | - | - |
|  | Cusp 7 | Coefficient | -0.01 | -0.02 | -0.07 | -0.37 | -0.46 | **-1.33** | -0.37 | -0.45 | **-1.21** | -0.08 | -0.10 | -0.31 | -0.07 | -0.09 | -0.30 |
|  |  | p-value | 0.68 | 0.46 | 0.08 | 0.35 | 0.19 | **0.03** | 0.30 | 0.15 | **0.02** | 0.49 | 0.31 | 0.06 | 0.52 | 0.33 | 0.24 |
|  |  | Odds ratio | - | - | - | - | - | **0.88** | - | - | **0.89** | - | **-** | - | - | **-** | **-** |
| RM_1_ | Cusp 5 | Coefficient | -0.03 | - | - | **-1.72** | - | - | -1.24 | - | - | -0.32 | - | - | -0.04 | - | - |
|  |  | p-value | 0.60 | - | - | **0.04** | - | - | 0.10 | - | - | 0.18 | - | - | 0.66 | - | - |
|  |  | Odds ratio | - | - | - | **0.84** | - | - | - | - | - | - | - | - | - | - | - |
|  | Cusp 6 | Coefficient | -0.00 | 0.01 | **-** | -0.22 | 0.20 | - | -0.15 | 0.19 | - | -0.03 | 0.07 | - | -0.04 | 0.04 | - |
|  |  | p-value | 0.89 | 0.59 | **-** | 0.55 | 0.54 | - | 0.65 | 0.51 | - | 0.79 | 0.48 | - | 0.66 | 0.65 | - |
|  |  | Odds ratio | - | - | - | - | - | - | - | - | - | - | - | - | - | - | - |
|  | Cusp 7 | Coefficient | 0.03 | 0.02 | -0.03 | 0.15 | 0.01 | -0.71 | 0.12 | -0.02 | -0.83 | 0.10 | 0.05 | -0.13 | 0.06 | 0.02 | -0.16 |
|  |  | p-value | 0.39 | 0.61 | 0.62 | 0.75 | 0.98 | 0.35 | 0.78 | 0.96 | 0.24 | 0.46 | 0.67 | 0.56 | 0.61 | 0.85 | 0.42 |
|  |  | Odds ratio | - | - | - | - | - | - | - | - | - | - | - | - | - | - | - |

**S18 Table. Results of logistic regression when using intercusp area as a predictor of lower molar accessory cusp development (morphology dichotomized).** Cells highlighted in blue represent intercusp areas that become significantly larger with accessory cusp development. Cells highlighted in red represent intercusp areas that become significantly smaller with accessory cusp development.

^1^α≤0.05; significant results in bold

^2^Odds ratio obtained by dividing the model coefficient by 10 to scale the odds ratio to a 0.10-unit change and exponentiating the quotient

^3^Absolute intercusp area formed by protoconid-metaconid-hypoconid-entoconid polygon

^4^Absolute intercusp area formed by protoconid-metaconid-hypoconid-entoconid-hypoconulid polygon

^5^Absolute intercusp area formed by protoconid-metaconid-hypoconid-entoconid-hypoconulid-cusp6 polygon

**S1 Figure. Data collection methodology of the current study.** Red circles mark the cusp tip location. The horizontal red line at the top of the images shows the linear size reference tool used to scale all measurements. The blue perpendicular lines indicate the mesiodistal and buccolingual tooth dimensions, which were collected following standards in the field [158-159]. The green polygon surrounding the crown demonstrates how the crown base was traced to measure its area. Image courtesy of senior author’s (KSP) personal collection.

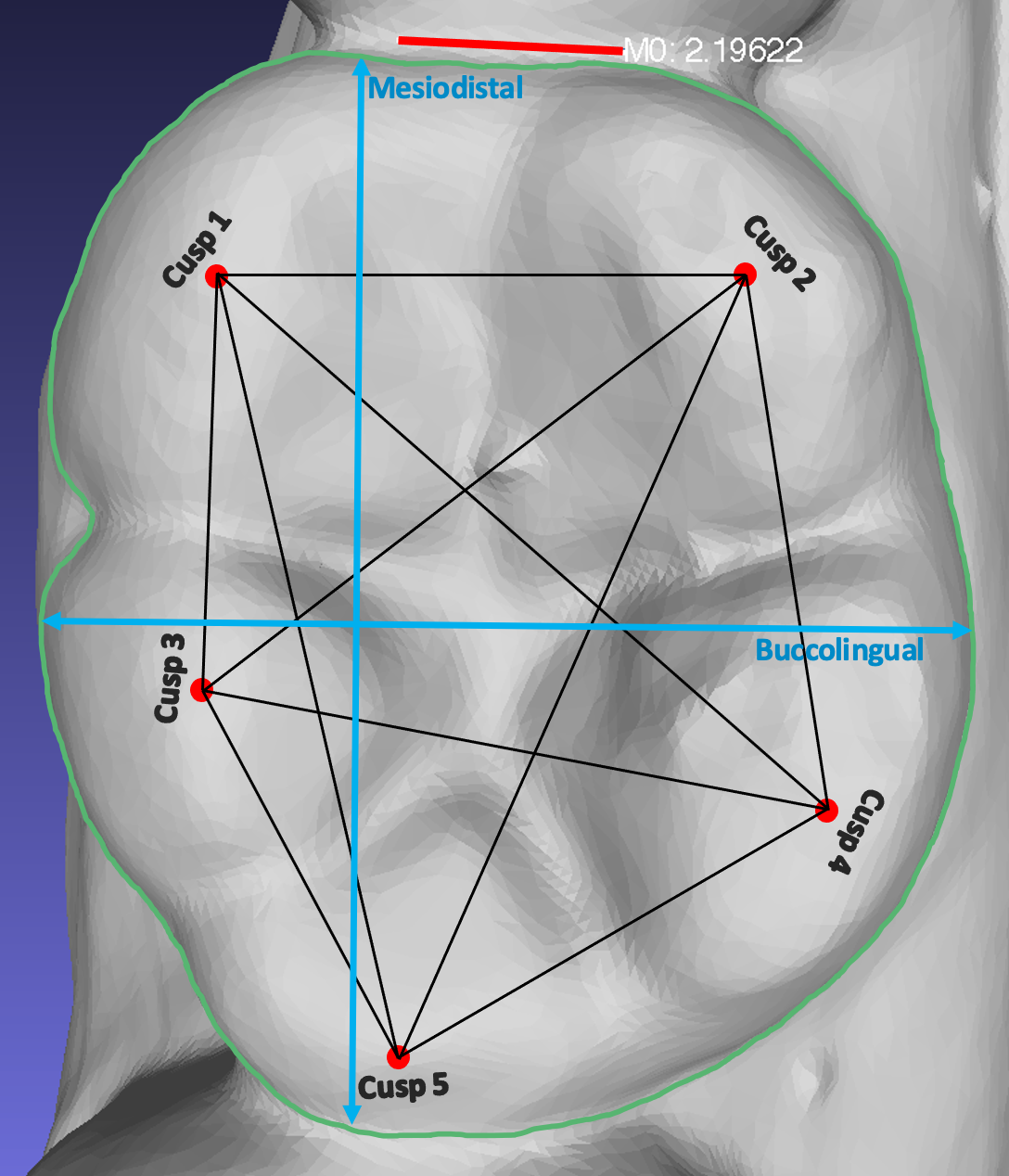

**S2 Figure. Relationship between cusp 5 trait expression and BLxMD crown area**

**S3 Figure.** **Relationship between cusp 6 trait expression and BLxMD crown area**

**S4 Figure. Relationship between cusp 7 trait expression and BLxMD crown area**

**S5 Figure.** **Relationship between cusp 5 development (dichotomized) and BLxMD crown area**

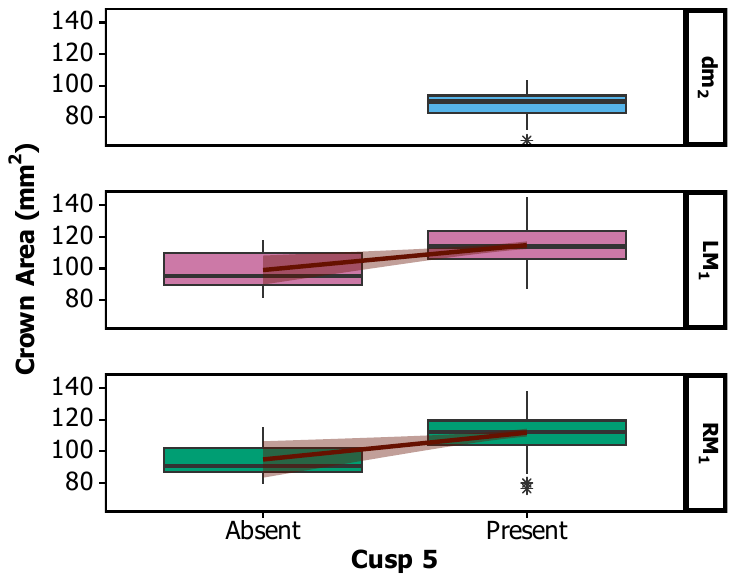

**S6 Figure.** **Relationship between cusp 6 development (dichotomized) and BLxMD crown area**

**S7 Figure.** **Relationship between cusp 7 development (dichotomized) and BLxMD crown area**

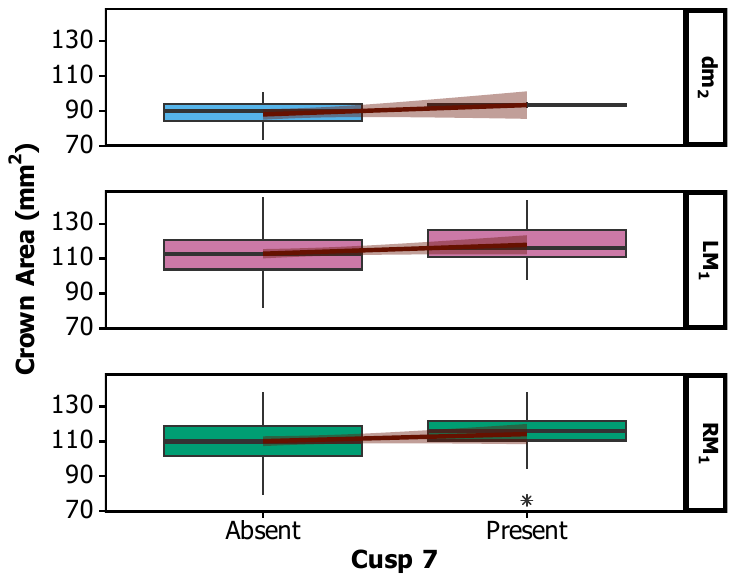

**S8 Figure.** **Relationship between cusp 5 trait expression and traced crown area**

**S9 Figure. Relationship between cusp 6 trait expression and traced crown area**

**S10 Figure.** **Relationship between cusp 7 trait expression and traced crown area**

**S11 Figure. Relationship between cusp 5 development (dichotomized) and traced crown area**

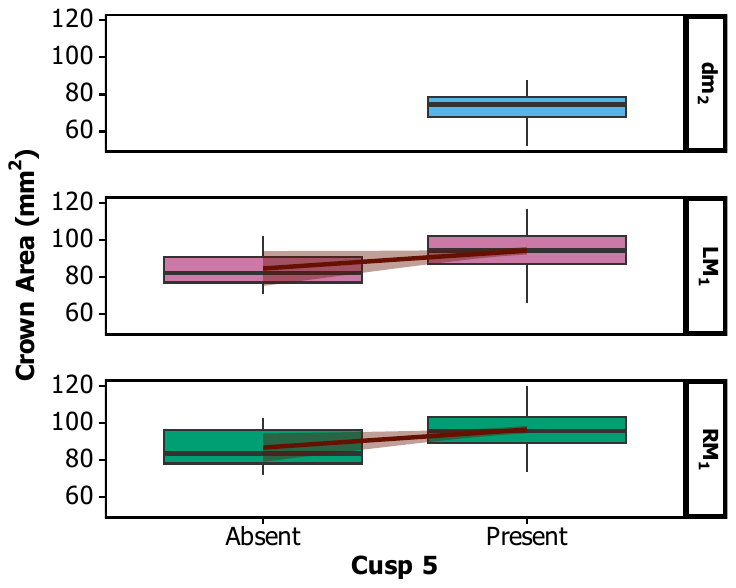

**S12 Figure. Relationship between cusp 6 development (dichotomized) and traced crown area**

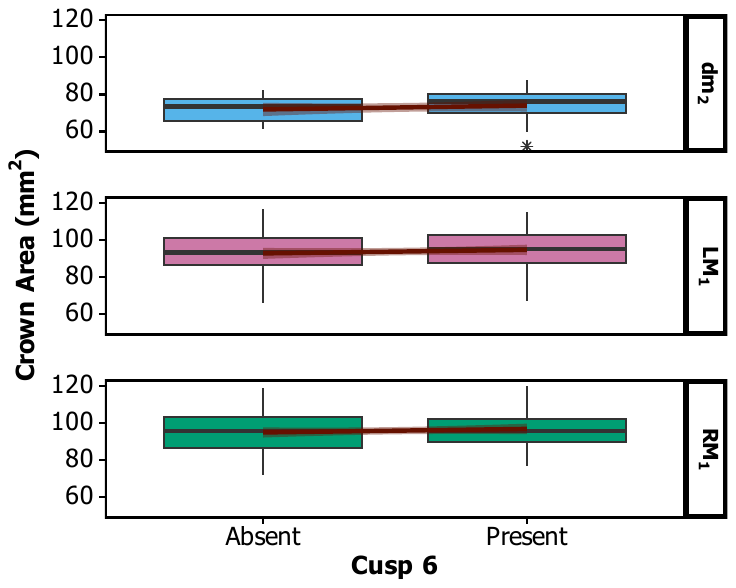

**S13 Figure. Relationship between cusp 7 development (dichotomized) and traced crown area**

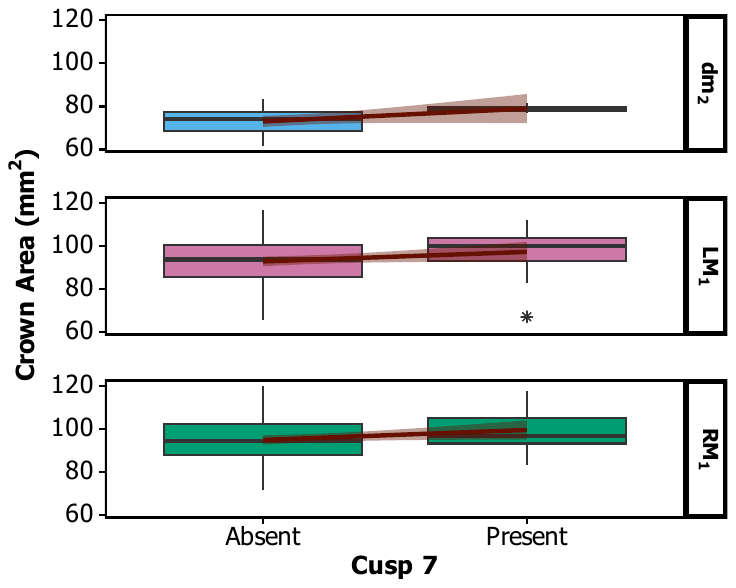

**S14 Figure.** **Relationship between cusp 5 trait expression and absolute intercusp distance**

**S15 Figure.** **Relationship between cusp 6 trait expression and absolute intercusp distance**

**S16 Figure. Relationship between cusp 7 trait expression and absolute intercusp distance**

**S17 Figure.** **Relationship between cusp 5 development (dichotomized) and absolute intercusp distance**

**S18 Figure. Relationship between cusp 6 development (dichotomized) and absolute intercusp distance**

**S19 Figure.** **Relationship between cusp 7 development (dichotomized) and absolute intercusp distance**

**S20 Figure.** **Relationship between cusp 5 trait expression and relative intercusp distance scaled by BLxMD crown area**

**S21 Figure.** **Relationship between cusp 6 trait expression and relative intercusp distance scaled by BLxMD crown area**

**S22 Figure.** **Relationship between cusp 7 trait expression and relative intercusp distance scaled by BLxMD crown area**

**S23 Figure.** **Relationship between cusp 5 development (dichotomized) and relative intercusp distance scaled by BLxMD crown area**

**S24 Figure. Relationship between cusp 6 development (dichotomized) and relative intercusp distance scaled by BLxMD crown area**

**S25 Figure.** **Relationship between cusp 7 development (dichotomized) and relative intercusp distance scaled by BLxMD crown area**

**S26 Figure. Relationship between cusp 5 trait expression and relative intercusp distance scaled by traced crown area**

**S27 Figure. Relationship between cusp 6 trait expression and relative intercusp distance scaled by traced crown area**

**S28 Figure. Relationship between cusp 7 trait expression and relative intercusp distance scaled by traced crown area**

**S29 Figure.** **Relationship between cusp 5 development (dichotomized) and relative intercusp distance scaled by traced crown area**

**S30 Figure.** **Relationship between cusp 6 development (dichotomized) and relative intercusp distance scaled by traced crown area**

**S31 Figure.** **Relationship between cusp 7 development (dichotomized) and relative intercusp distance scaled by traced crown area**

**S32 Figure. Relationship between cusp 5 trait expression and relative intercusp distance scaled by mesiodistal (MD) tooth dimension**

**S33 Figure. Relationship between cusp 6 trait expression and relative intercusp distance scaled by mesiodistal (MD) tooth dimension**

**S34 Figure.** **Relationship between cusp 7 trait expression and relative intercusp distance scaled by mesiodistal (MD) tooth dimension**

**S35 Figure.** **Relationship between cusp 5 development (dichotomized) and relative intercusp distance scaled by mesiodistal (MD) tooth dimension**

**S36 Figure.** **Relationship between cusp 6 development (dichotomized) and relative intercusp distance scaled by mesiodistal (MD) tooth dimension**

**S37 Figure.** **Relationship between cusp 7 development (dichotomized) and relative intercusp distance scaled by mesiodistal (MD) tooth dimension**

**S38 Figure.** **Relationship between cusp 5 trait expression and relative intercusp distance scaled by buccolingual (BL) tooth dimension**

**S39 Figure.** **Relationship between cusp 6 trait expression and relative intercusp distance scaled by buccolingual (BL) tooth dimension**

**S40 Figure.** **Relationship between cusp 7 trait expression and relative intercusp distance scaled by buccolingual (BL) tooth dimension**

**S41 Figure.** **Relationship between cusp 5 development (dichotomized) and relative intercusp distance scaled by buccolingual (BL) tooth dimension**

**S42 Figure.** **Relationship between cusp 6 development (dichotomized) and relative intercusp distance scaled by buccolingual (BL) tooth dimension**

**S43 Figure.** **Relationship between cusp 7 development (dichotomized) and relative intercusp distance scaled by buccolingual (BL) tooth dimension**

**S44 Figure.** **Relationship between cusp 5 trait expression and intercusp area**

**S45 Figure.** **Relationship between cusp 6 trait expression and intercusp area**

**S46 Figure.** **Relationship between cusp 7 trait expression and intercusp area**

**S47 Figure.** **Relationship between cusp 5 development (dichotomized) and intercusp area**

**S48 Figure. Relationship between cusp 6 development (dichotomized) and intercusp area**

**S49 Figure.** **Relationship between cusp 7 development (dichotomized) and intercusp area**
