## Supplementary material for "Evaluating the patterning cascade model of tooth morphogenesis in the human lower mixed and permanent dentition": S2 Appendix

**Materials and Methods**

**Study Sample**

The study sample includes digital images of modern human dental casts collected from Solomon Islander individuals during the course of the Harvard Solomon Islands Project (HSIP) [162]. The HSIP collected dental casts from individuals belonging to several language groups, most of which are currently housed at the Peabody Museum of Archaeology and Ethnology at Harvard University. In collaboration with the Peabody Museum, the Dental Phenomics Project has generated anonymized scans and images of participant dentitions, which were utilized as part of this study. The sample includes 124 individuals from the island of Malaita for whom dm_2_-M_1_ and/or RM_1_-LM_1_ pairs were present in the dental arcade, cusp tip placement was clearly discernible, and dental morphological trait expression could be determined. No personally identifying information was accessed as part of this research. This study was exempted from review by the University of Arkansas’ Office of Research Integrity and Compliance and the Institutional Review Board of Arizona State University (see Ethics Statement).

**Data Collection**

Dental data were collected from digital 3D scans of anonymized mandibular dental casts capturing the mixed or permanent dentition. All data were collected by the primary author (DEK), except deciduous morphology, which was collected by the senior author (KSP). Individuals were included in the study sample if the entire crown under observation had completed gingival eruption, secondary enamel knot location could readily be estimated based on cusp tip placement for all cusps, and morphological trait scores could be assessed based on cusp shape and size. Individuals with excessive cusp wear obliterating cusp tip location and accessory cusp morphology, as well as casts exhibiting casting errors (i.e., bubbles, nodules) were excluded from the study. Whenever observable, the dm_2_-M_1_ metamere and the RM_1_-LM_1_ antimere pair were examined. Due to extensive dm_2_ wear and sample size concerns, examination of deciduous antimeres was not possible. For metameric comparisons in the mixed dentition, the dm_2_ least affected by dental wear was selected for analysis and the permanent M_1_ located on the same side as the dm_2_ was examined. Whenever the M_1_ on the same side as the dm_2_ could not be observed, values from its antimere were substituted. The data collection and analysis protocols employed in the current study closely follow those utilized by Hunter and colleagues [128], Moormann and colleagues [135], and Paul and colleagues [134].

The 3D scans of mandibular casts were visualized in the 3D mesh processing software system MeshLab v2022.02 [163]. Cusp tips were marked for all cusps visible on the crown surface and the measuring tool was used to create a linear size reference in millimeters just above the crown area. Including the size reference, a screenshot was taken of the crown from an occlusal view. This image was transferred to the Java-based image processing software ImageJ v.1.53, where intercusp distance and crown area data were collected [164]. Using the linear size reference included on the image, all measurements were calibrated to scale. Intercusp distances were collected between all cusps present on the crown using the line selection tool (S1 Figure). We utilized crown base area as a proxy corresponding to developmental timeframe since previous studies found larger crowns to be characterized by extended periods of gestation [165-166]. Crown area was measured in two ways: by tracing base area using the polygon selection tool (trace area) and by multiplying the 2D linear measurements of maximum buccolingual (BL) and mesiodistal (MD) diameter (BLxMD).

Dental morphological data were collected for LM_1_ and RM_1_ using the 3D digital scans visualized in MeshLab. For dm_2_, morphology data were collected through primary observation. A previous precision study using casted dentitions showed high correspondence in resulting morphology datasets when comparing scores recorded from 3D scans to those collected via primary observation; mean error for lower accessory cusps ranged from 0.04-0.09 grade and percent concordance ranged from ~81-93% [25]. Data collection and dichotomization protocol for the permanent molars followed the Arizona State University Dental Anthropology System (ASUDAS) (S1 Table) [160-161]. The ASUDAS quantifies dental morphological variation on an ordinal scale to approximate the underlying continuous genotypic distribution that culminates in the phenotype [167-168]. Morphological data were collected for cusp 5 (hypoconulid), cusp 6 (*tuberculum sextum*), and cusp 7 (*tuberculum intermedium*) (Figure 2). The “1A” grade of the cusp 7 scoring system was collected from the images; however, this manifestation corresponds to a shouldering of the metaconid, representing a *metastylid* rather demarcating a true accessory cusp that emerges from the mesial ridge between the metaconid and entoconid (grades 1-4) [153]. The emergence of the “metastylid-type” cusp 7 precedes emergence of the “interconulid-type” grades, further supporting the idea that these are developmentally distinct structures [89]. Therefore, individuals exhibiting a 1A grade of cusp 7 were eliminated from statistical analyses associated with cusp 7. While the ASUDAS is designed to capture morphological trait variation in the permanent dentition, this system has been successfully applied to the deciduous dentition and was utilized in this study [25, 146, 152, 169].

**Statistical Methods**

All statistical analyses were conducted in RStudio v4.2.2 [170-171]. Inter- and intra-observer error was determined for a small subset of the original sample (15 individuals). DEK re-measured dimensions and re-scored morphological trait expression for permanent teeth approximately six months after initial data collection to establish intra-observer error rates. Inter-observer agreement was calculated for morphological data by comparing trait scores between DEK and CMS (cusp 6 and 7) and KSP (cusp 5) for the permanent dentition. Error rates for deciduous morphology have been consistently within an acceptable range [25, 27, 134, 146, 152]. Cohen’s Weighted Kappa was utilized to identify agreement rates for dental morphological trait scores using the *irr* package [172]. This statistic highlights rater agreement by capturing the magnitude of score differences between raters/scoring sessions, weighing large disparities more heavily. Mean absolute error, mean absolute percentage error, and root-mean-square error rates were determined to evaluate intra-observer error for the continuous data using the *Metrics* package [173]. Technical error of measurement (TEM) and relative technical error of measurement (%TEM) were also calculated for the metric variables [174-176]. Beyond establishing measurement error, paired tests were applied to all metric data collected for the same variable at two different times to determine if discrepancies were significant (α<0.05).

Prior to statistical analysis, Shapiro-Wilk tests were run for all variables tested in metameric (dm_2_-M_1_) and antimeric (RM_1_-LM_1_) comparative analyses to evaluate if the continuous data were normally distributed. Paired t-tests were used whenever results of the Shapiro-Wilks tests were non-significant, while paired-sample Wilcoxon signed-rank tests were applied to all ordinal morphological data and metric data with a non-normal distribution. Relative intercusp distances were calculated to eliminate crown size as a confounding variable by dividing the absolute intercusp distance (AICD) by the square root of the crown area (SQRTA) (*i.e.,* RICD = AICD/SQRTA). Absolute intercusp area (AICA) data were collected in ImageJ by measuring the polygonal area created by the cusp tips and scaled to a relative value to eliminate tooth size as a confounding variable (RICA).

Using the *MASS* package, proportional odds logistic regression was performed to determine if the expression of later-developing accessory cusps (*i.e.*, cusp 5, cusp 6, and cusp 7) (dependent variable) was influenced by the distance between earlier-developing cusps (AICD or RICD), tooth size (BLxMD or trace area), or intercusp area (AICA or RICA) as predicted by the patterning cascade model of tooth morphogenesis [177]. The same relationships were examined using dichotomized morphological trait scores with grade 1 as the breakpoint for all morphological traits [160-161]. Because the dichotomized data only has two levels (*i.e.,* trait presence/absence), logistic regression was employed to establish if the development (rather than expression) of accessory cusps was influenced by any of the PCM-predicted factors. Brant tests were conducted using the poTEST function from the *MASS* package to establish if the proportional odds regression models met the assumption of proportionality [177-178]. The assumption of proportionality in the context of the current study implies that the coefficients describing the relationship between levels of the response variable are equal and their regression lines are parallel. Using the *car* package, likelihood ratio chi-square tests were calculated to determine the significance of model fit (α≤0.05) [179]. Odds ratios were calculated to ascertain the degree to which the independent variable of interest (intercusp distance, tooth size, or intercusp area) impacts the likelihood of forming an accessory cusp with more pronounced expression. This was achieved by dividing the model coefficient by 10 to scale the odds ratio to a 0.1-unit change and then exponentiating the quotient.
